# GAIT-GM: Galaxy tools for modeling metabolite changes as a function of gene expression

**DOI:** 10.1101/2020.12.25.424407

**Authors:** Lauren M. McIntyre, Francisco Huertas, Olexander Moskalenko, Marta Llansola, Vicente Felipo, Alison M. Morse, Ana Conesa

**Affiliations:** Department of Molecular Genetics and Microbiology, University of Florida, Gainesville Florida, 32610; UF Genetics Institute, University of Florida, Gainesville Florida, 32608; Department of Microbiology and Cell Science, University of Florida, Gainesville Florida, 32611; UF Research Computing. University of Florida, Gainesville Florida, 32610; Prince Felipe Research Center, Avda. Eduardo Primo Yúfera 3, 46012, Valencia, Spain

**Keywords:** Galaxy, data integration, transcriptomics, metabolomics

## Abstract

Galaxy is a user-friendly platform with a strong development community and a rich set of tools for omics data analysis. While multi-omics experiments are becoming popular, tools for multi-omics data analysis are poorly represented in this platform. Here we present GAIT-GM, a set of new Galaxy tools for integrative analysis of gene expression and metabolomics data. In the Annotation Tool, features are mapped to KEGG pathway using a text mining approach to increase the number of mapped metabolites. Several interconnected databases are used to maximally map gene IDs across species. In the Integration Tool, changes in metabolite levels are modelled as a function of gene expression in a flexible manner. Both unbiased exploration of relationships between genes and metabolites and biologically informed models based on pathway data are enabled. The GAIT-GM tools are freely available at https://github.com/SECIMTools/gait-gm.

## Introduction

High-throughput transcriptomics and metabolomics are now common analytical techniques in many labs, where scientists not expert in bioinformatics or statistics demand user-friendly tools for data analysis. The Galaxy platform is a widely used open source community development with an easy to understand GUI and a ‘mix and match’ pipeline building philosophy, that has become one of the most successful resources for omics data analysis by biologists. Galaxy contains tools to analyze gene expression and for untargeted metabolomics^1,2^. However, Galaxy tools for joint analysis of transcriptomics and metabolomics are scarce^3^ and this limits the utilisation of this platform to address multi-omics data integration problems.

A diversity of methods have been developed that integrate metabolomics and gene expression data^4–7^. Many of the existing integration approaches are based on multivariate dimension reduction techniques that assess the covariation structures between transcript and metabolite levels^8,9^. However, these methods rarely incorporate existing biological knowledge, which hampers the interpretation of analysis results. Pathway database such as KEGG^10^, that include both metabolites and genes, can be used to map features to a common biological process and to guide integrative analysis^11^. However, this strategy has also limitations. Mapping measured metabolite levels to a pathway database such as KEGG is restricted to the metabolites that are actually annotated to metabolic pathways. Many detected metabolites, especially lipids, cannot be mapped to the pathway data, and are eventually left aside in the integrative analysis. Moreover, despite the clear evidence of the importance of using metabolite identifiers^12^, and the existence of several metabolite ID conventions such as InChIKey and PubChemID^13^, natural language is still prevalent, hampering the use of databases that rely on identifiers. At present, common natural language names are not standardized in the chemical community. Moreover, using names for metabolites introduces variability due to different technology providers using different conventions. For example, using *lactate* versus *lactic acid* (the conjugate base of an acid versus the acid) or using *α-ketoglutarate* versus *2-oxoglutarate*. Moreover, metabolomics platforms usually do not resolve isomeric variants of sugars such as L-glucose and D-glucose, for which a different notation exists in KEGG. Finally, existing pathway tools^14–17^ have limited or no choices for the pre-processing of gene expression and metabolomics data, and typically do not incorporate methods to identify significant differences across experimental conditions. The result is that each scientist must deploy several additional tools for a successful and complete analysis. Using a series of different tools requires several data transfers and reformats, and the analysis can become fragmented and hard to reproduce. In contrast, linking the same tools inside Galaxy results in an integrated and documented workflow that is immediately reproducible, without extra documentation. Here we present Galaxy tools that integrate gene expression and metabolites by mapping to KEGG pathways and use a text-mining algorithm to improve metabolites identification. We hypothesize that metabolite changes are a function of transcriptional regulation and have created a flexible Integration Tool to build models of gene-metabolite regulation in both a biologically based and an unbiased mode. Leveraging the Galaxy framework, pipelines with pre-processing steps for differential expression of genes and metabolites can be easily created and saved for future studies.

## Results

### General overview of GAIT-GM

GAIT-GM (Galaxy Annotation and Integration Tools for Genes and Metabolites) is a set of tools for the analysis of metabolomics and transcriptomics data that enable the development of Galaxy workflows for statistical integration of gene and metabolite expression (**Figure 1**). Starting from a paired metabolomics and transcriptomics dataset, the Annotation Tools efficiently map gene IDs and metabolite names to KEGG IDs and KEGG Pathways. Features can be filtered using existing differential expression analysis tools to create a reduced dataset of significant features to use in the integrative analysis. The Integration Tools includes a diversity of analysis options to correlate genes and metabolites either based on the data alone or using pathway information to guide the analysis. GAIT-GM returns networks, text files and heatmap graphs as output (**Figure 1A**).

**Figure 1.**
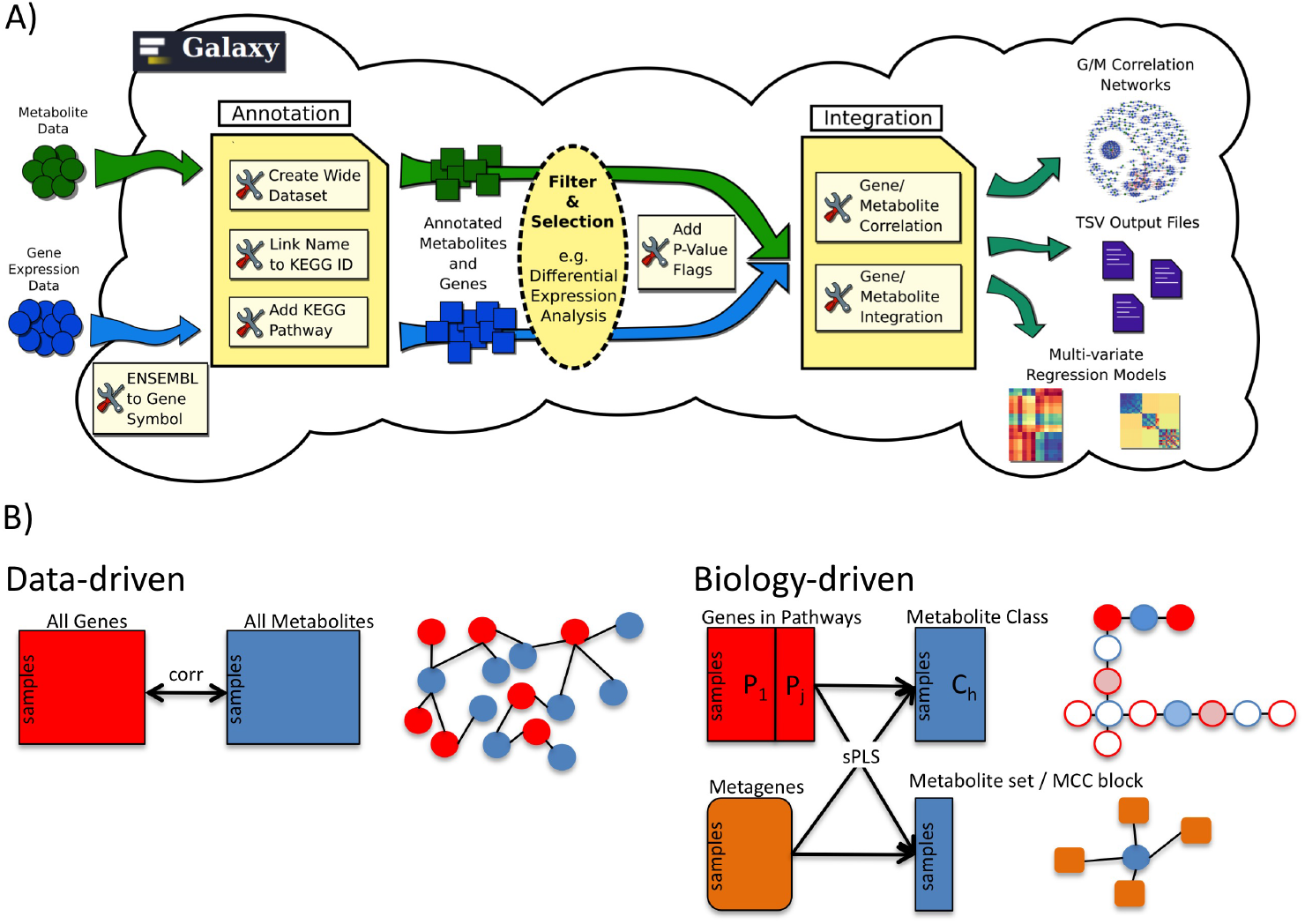
Galaxy pipeline for transcriptomics-metabolomics integration enabled by GAIT-GM. A) Annotation and Integration Tools are provided by GAIT-GM. Filter and Selection tools are taken from existing Galaxy tools. B) Analysis flexibility of the GAIT-GM tool. Unbiased -based on a correlation metric- and biologically informed -based on pathway, metabolite class and Partial Least Square analysis-analyses are implemented.

The GAIT-GM framework for the integration of metabolomics with gene expression data builds on four biological hypotheses and the principle of analysis flexibility (**Figure 1B**). The first hypothesis is that gene expression (indirectly) regulates metabolite levels and this regulatory relationship can be identified as a relationship between the quantitative levels of the metabolite, and the transcript. This principle underlies both the unbiased analyses and the biologically informed analyses. The second hypothesis is that genes/metabolites belonging to the same pathway are more likely to be engaged in regulatory relationships. In the biologically informed analyses, shared Pathway information is used to develop models of metabolite level changes. The third hypothesis is that metabolites that belong to the same “class” (i.e. all sphingomyelins) may have a common underlying regulator. For the unbiased approach, co-variation patterns are used to cluster metabolites. GAIT-GM implements tools that group metabolites to allow joint analysis in both unbiased and biologically informed ways. Fourth, pathways as a whole can be considered functional units regulating metabolic changes. GAIT-GM enables the estimation of a pathway effect. Using the principle of flexibility each tool is developed in a modular format, allowing tools to be chained together into a pipeline or to enable different tools to be used for genes and metabolites. Multiple combinations or subsets of genes and metabolites can be integrated to answer different research questions.

### Application to a mouse dataset

We demonstrate the utilization of the GAIT-GM tools using a rat experiment that evaluates the effects of high ammonium acetate levels in Wilstar rats. Rats were fed during 4 weeks with a 25% of ammonium acetate to induce Hyperamonemia or a normal diet (Control). After this period of time, all rats were sacrificed cerebellum was isolated. One half was used for RNA extraction and the other half was immediately frozen to subsequent metabolomics analysis. 50 bp single read RNA-seq run with Illumina 2500 Hiseq. Data were processed as described^18^ and a gene expression dataset was obtained with 13,013 genes. Metabolomics profiles were obtained with a BIOCRATES instrument for LC-MS. A total of 131 compounds were measured.

#### GAIT-GM Annotation Tool improves feature mapping to pathways

GAIT-GM implements a number of parsing, text mining and database cross-reference strategies to maximally map gene ID and metabolite names to the KEGG database. For our rat dataset this approach increased the mapping of user metabolites to KEGG compound IDs from 26% to 95 % and most of them could also be located to at least one Pathway (**Figure 2A**). This significant improvement is partly achieved by the flexible parsing of metabolite names (**Figure 2C**), but also by the assignment of structural lipids (such as sphingomyelins, phosphatidylcholines, ceramides, etc) with multiple compounds of variable side chain lengths to a generic lipid class present in the KEGG database (**Figure 2D**). The results in lipids having on average 30 compounds per KEGG ID, while other metabolites have a 1:1 relationship to KEGG IDs (**Figure 2E**). While, most genes were mapped to KEEG IDs, only 20% mapped to pathways (**Figure 2B**).

**Figure 2.**
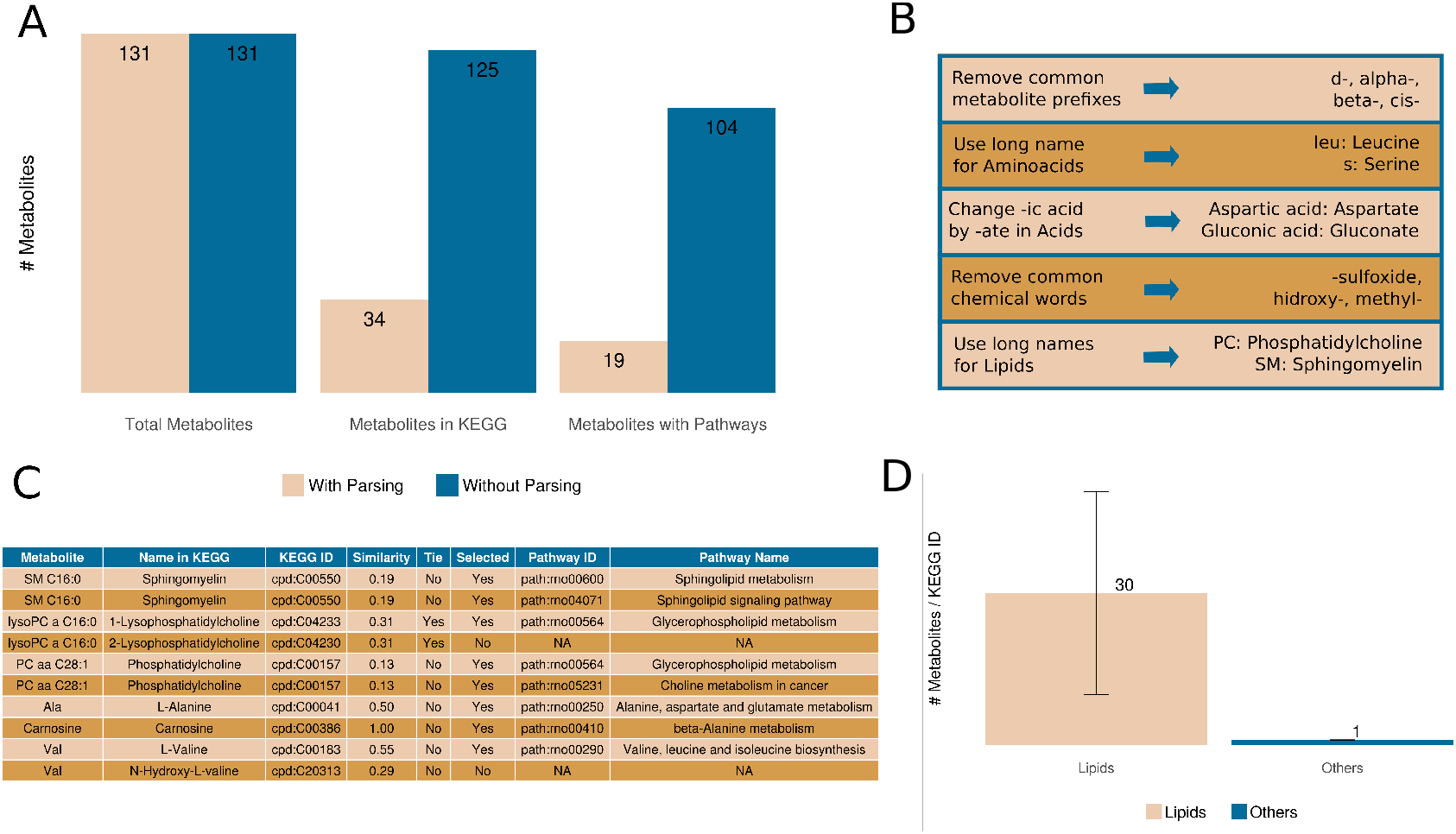
GAIT-GM improvement of mapping of features to KEGG database. A) Mapping improvement at metabolites. B) Mapping results for GeneIDs C) Example of text-mining rules. D) Example of mapping output for metabolites. C) Cardinality of input metabolites to KEEG compound IDs as a function of the metabolite type.

#### GAIT-GM Integrative Tools characterise the gene-metabolite co-variation network

The GAIT-GM Integration module, can estimate all possible gene-metabolite correlations (Pearson, Spearman or Kendall). Analysis of the top 500 gene-metabolite correlations in the rat dataset, reveals a network composed of multiple small modules with one central metabolite linked to a few gene ID, and two larger connected components, enriched in structural lipids (Supplementary Figure 1). Interestingly, a prominent network component contains one Phosphatidylcholine (PC aa 30:0) that is highly correlated with 57 genes, that are enriched for intracellular membrane and transport functions, functions associated to PC lipids cellular roles^19^. These results suggest that further analysis of co-expression patterns of these lipids might be meaningful to unravel putative functional or regulatory relationships with expressed genes.

#### GAIT-GM biologically-informed integration reveals regulatory differences for structural lipids

The sphingomyelin component of the rat dataset is composed of twelve separate metabolites all identified as sphingomyelins. We ask whether all metabolites in this class behave similarly or whether subsets exist regulated by different genes. The GAIT-GM integration tool was used to build an sPLS model^20^, where the response matrix is the set of twelve sphingomyelins, and the predictor matrix is composed of differentially expressed genes in KEGG pathways containing at least one sphingomyelin. We identified two subsets of sphingomyelins differentially regulated by four sets of genes (**Figure 3**). Notably, one pattern corresponds to long chain and the other to short chain compounds. Mapk9, involved in sphingomyelin signalling, was associated with long chains, while Sptlc2, a key enzyme in sphingomyelin biosynthesis, was related to short chains. The results suggest that gene regulation of sphingomyelins may differ based on the size and function of the lipid chain.

**Figure 3.**
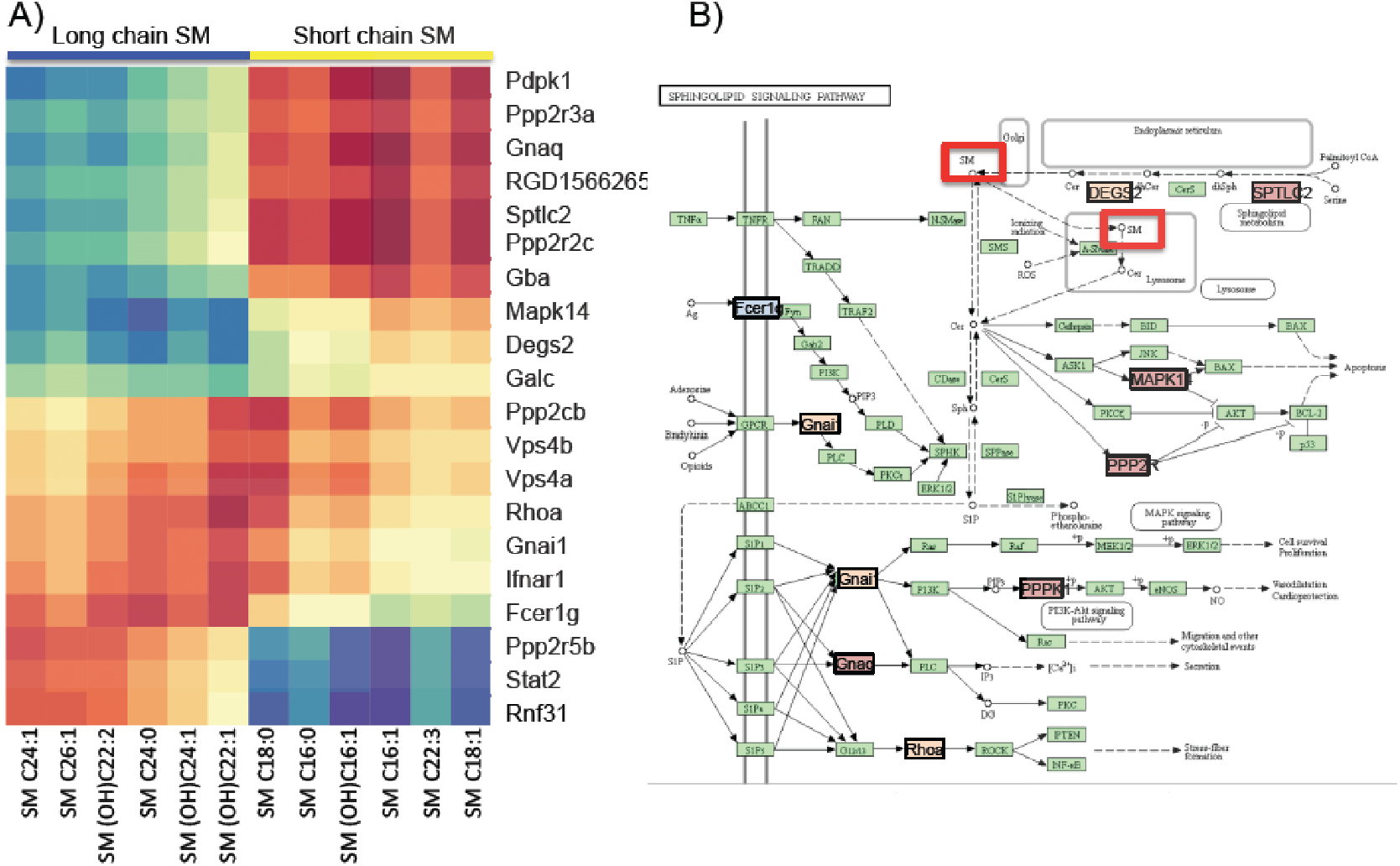
GAIT-GM integrative analysis. A) sPLS model of sphingomyelins vs. pathway genes. B) Sphingolipid signaling pathway localization of significant genes.

#### GAIT-GM Integration Tools identify novel functional relationships at the pathway level

We hypothesized that global transcriptional patterns may influence global metabolic changes. To explore this idea, we used MMC to cluster metabolites into modules based on co-expression^21^ and computed pathway metagenes^22^. MCC identified ten metabolite modules (Supplementary Figure 2). Metabolites in Modules 1 and 2 are positively correlated and represent mostly long chain phosphatidylcholines (PC). Metabolites in Modules 3 and 4 are negatively correlated to 1 and 2 and contain mostly short chain PC. Metagenes from all pathways containing at least 3 genes were used to build an sPLS models for these 4 MMC metabolite modules (**Figure 4**). We identified several metabolic and signalling pathways associated with the MMC modules. For example, *Butanoate metabolism* –involved in the creation of lipid precursors-, *Glutathione metabolism* –relevant in detoxification-, and *NFKB and Natural Killer Mediated Toxicity* were positively correlated with Modules 1-2, while *Insulin secretion* and *Inositol Phosphate metabolism* were associated with Modules 3-4. These results suggest different regulatory and functional specificities for different subsets of the PCs.

**Figure 4.**
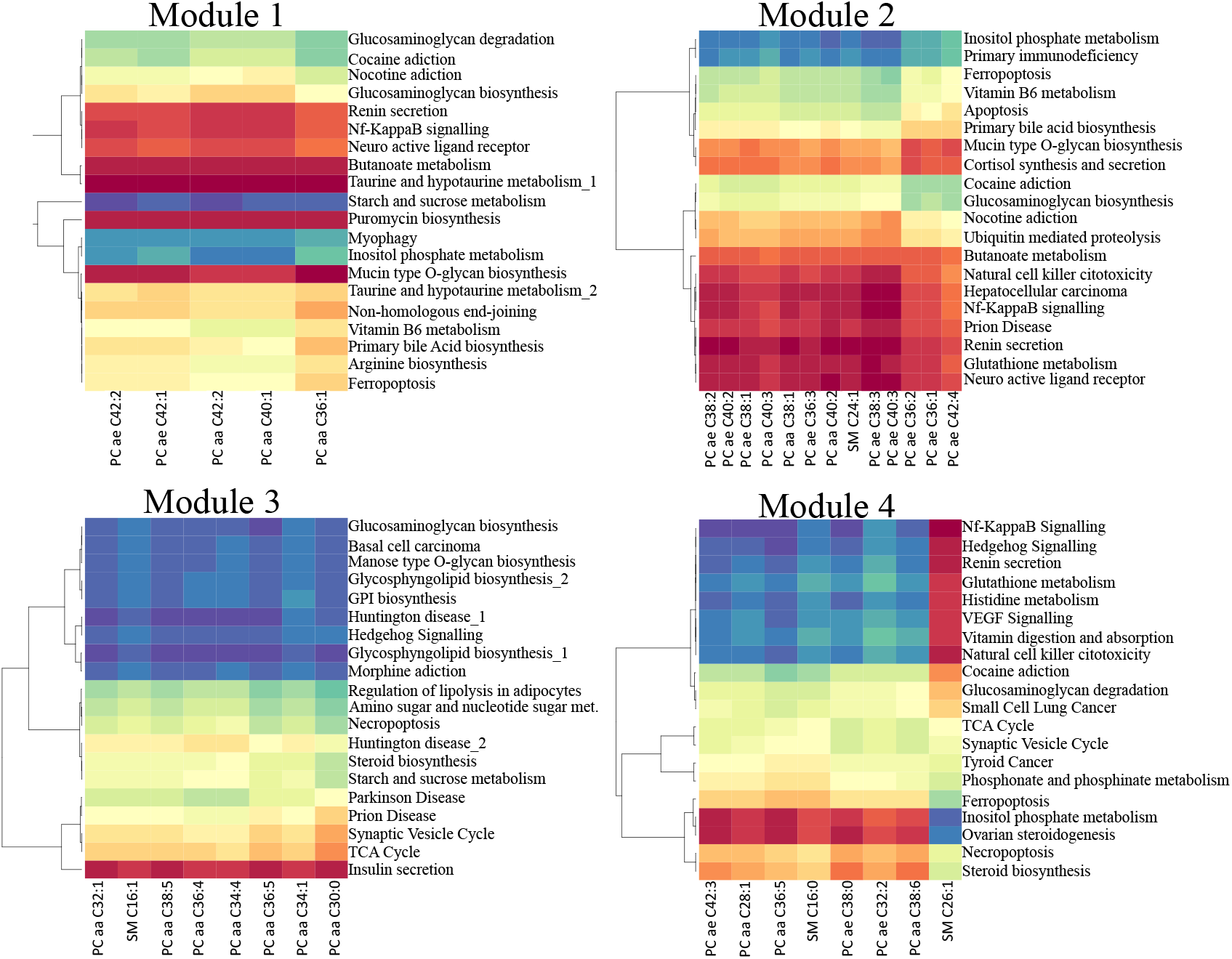
GAIT-GM Integrative Tool analysis results. sPLS analysis of 4 MCC metabolite modules versus pathway metagenes. Each heatmap represents the correlation of a lipid clusters to the most coordinated gene expression signature of each pathway. Colour scale indicates high positive correlation as red and high negative correlation as blue.

## Discussion

Multiomics experiments including gene expression and metabolomics data are now within the reach of many biological research laboratories that do their own data analysis. Traditionally, such labs have used user-friendly bioinformatics tools that allow easy access to state-of-the art algorithms and make possible interactive and exploratory analysis of the data. Galaxy is a platform widely used in these cases because it allows flexible configuration of analysis pipelines to accommodate a wide range of experimental designs and analysis needs. However, not many tools for integrative transcriptomics and metabolomics exist in Galaxy. The GAIT-GM fills a gap in multiomics data analysis for the Galaxy project.

There are several types of strategies for integrating gene expression and metabolomics data, and each of them pose different bioinformatics needs. When *a priori* information on the identity and functional relationship between genes / metabolites is not known or not incorporated in analysis, data can be modelled in *unbiased* ways. A straightforward approach is to search for correlation between genes and metabolites and look for patterns in results *post hoc*. This is implemented in the GAIT-GM Correlation module and can be used for exploratory unbiased analysis of the entire data set. This can be applied to all data, or only to genes/metabolites that are differentially expressed according to a factor in the experimental design (recommended). Alternatively, metabolites/genes can be grouped by co-expression using an unbiased clustering, the resulting groups can be modelled and/or the group can be further reduced to a single variable. For example suppose a co-variation analysis identifies three groups of metabolites and 5 groups of genes, an sPLS on the individual metabolites and genes for all group combinations can be performed (15 combinations in the above example). Another option is to reduce the gene expression patterns into 5 variables, one for each group and then for each group of metabolites, to model the 5 groups of co-expressed genes. These possibilities are implemented in the GAIT-GM Integration tool.

When biological knowledge is present, a more informed approach is possible and Tools in GAIT-GM have been developed to facilitate this approach. One important aspect is the annotation of metabolites. If compound IDs are present, a general tool links these IDs to the KEGG database and KEGG pathways. If compound IDs, are not present, or not uniformly present, the GAIT-GM Annotation Tool applied a number of text mining and parsing strategies based on the compound name that significantly improve the mapping of compounds of KEGG pathways. Another problem is the high number of lipidic compounds that have not a precise match in the database and for which a specific location in a metabolic pathway is simply unknown. These are the many phosphatidylcholines, sphingomyelines, ceramides, etc., that populate metabolomics datasets with varying lengths and saturation of their side chains. These compounds, apart from lacking a precise match in the KEGG database, may present high heterogeneity in their quantities and variation patterns across samples, making their analysis difficult. The GAIT-GM Tool has been specially designed to address this problem by allowing selection and clustering of metabolite classes thereby enabling the identification of different subsets of lipids of the same generic class. These subsets can then be modelled separately. In the rat dataset, we showed that the separation of lipid classes identifies putative distinct functional roles for different metabolite subsets.

In summary, the GAIT-GM provides a user-friendly analytical framework to integrate metabolomics and gene expression information. GAIT-GM is modular and allows a flexible integration that enables everything from a completely unbiased analysis to a pathway-centric biologically informed analysis, and various combinations of these integrative strategies. The Galaxy platform enables the construction of reproducible pipelines and transparent sharing of analytical approaches and results. There are two important yet unmet needs in the analysis of metabolomics data specifically developed here: the efficient mapping of compound names to KEGG, and the analysis of the heterogeneity and co-regulation with gene expression to improve their functional characterisation.

## Materials and methods

There are three basic steps for an integrated expression and metabolite analysis 1) Feature annotation 2) Feature selection 3) Integrated analysis. We have developed Galaxy tools that cover the first and third steps of this process and leverage existing Galaxy tools for the second step. The GAIT-GM Annotation Tools map common metabolite and gene names to KEGG IDs and associate them their common pathways. The GAIT-GM Integration Tools implement methods for unbiased (data-driven) and biologically informed integration for genes and metabolites. GAIT-GM Tools are wrapped for Galaxy and deposited in PyPi (https://pypi.org/project/gait-gm/) with a corresponding Conda recipe to enable users access on the command line. All code, including the needed Galaxy wrappers is also available on our github (https://github.com/SECIMTools/gait-gm).

Here we describe the methods behind GAIT-GM tools. We also provide examples of workflows that demonstrate several data integration options. Further technical details on tools utilisation are provided in the Supplementary Material User Guide.

### Annotation

#### Resolving metabolite names

To address the problem of matching of natural language metabolite names, typically provided by metabolomics facilities, to KEEG, GAIT-GM maps names onto KEGG IDs using a set of rules for the processing of natural language. The set of rules in PaintOmics3^11^ was used and expanded to assign metabolite names to the most likely compound in the KEGG database. For each input metabolite, a list of potentially related metabolites based on the similarities in their names is generated as follows: 1) Metabolite names are parsed according to the rules listed in the User Guide provide as Supplementary Material. Common metabolite prefixes are removed (cis-, trans-, d-, l-, (s)-, alpha-, beta-, alpha, beta, alpha-d-, beta-d-, alpha-l-, beta-l-, l-beta-, l-alpha-, d-beta-, d-alpha-). If the metabolite name given is an acid, then the name is modified to the conjugate base by replacing “ic acid”, “icacid” or “ic_acid” with “ate”. If amino acids are given in 1-letter or 3-letter abbreviations, names are modified to the full amino acid name. The following commonly used lipid abbreviations are modified to reflect the full names (SM = sphingomyelin, lysopc = lysophosphatidylcholine, PC = phosphatidylcholine, PE = phosphatidylethanolamine and LysoPE = lysophosphatidylethanolamine). Similarly, abbreviations for other commonly assayed metabolites are modified to reflect the full names (cit = citrate, orn = ornithine, thyr = thyroxine and boc = butoxycarbonyl). 2) Names are matched to KEGG. 3) A similarity score is calculated using the python internal SequenceMatcher class from difflib (https://docs.python.org/2/_sources/library/difflib.rst.txt, module and section author Tim Peters) that returns a measure of the similarity between two strings. Similarity score is based on the longest contiguous matching subsequence that does not contain ‘junk’ elements where ‘junk’ elements are defined as duplicates making up more than 1% of a sequence with minimum length of 200. Identical names receive a score of 1. 4) The highest similarity score is selected. 5) When the best match is tied with at least one other compound in KEGG, all matches are returned. A tie is determined if the similarity score is greater than 95% for 2 or more matches in the metabolite name. In this cae, a default selection is provided, but other possibilities are visible to the scientist and the selection can be easily modified before the next step.

#### Resolving gene names

Gene ID conversion has been a long and general problem of genomics databases. Tools such as DAVID^23^ and BridgeDB^24^ address this problem, although limitations exist, such as the number of species covered or the number of items that can be processed at a time. The KEGG Mapper is inconsistent in its naming conventions for different species. To our knowledge there is no general tool that links KEGG IDs for all species. We have adopted and improved the PaintOmics3 procedure for gene mapping^11^. Basically, PaintOmics3 fetches the ID translation information from public databases such as Ensembl, PDB, NCBI Refseq and KEGG, generates the translation tables and stores them in MongoDB collections. For example, given a feature ID (gene, protein or transcript) for database A, to translate to a valid gene name for database B, first the system retrieves the list of transcripts associated with the feature (if any). Then, for each transcript ID in database A, the equivalent transcript identifier at database B. Finally, to translate back to genes, the system finds the gene name associated to each identified transcript. Although this method has some limitations, mainly due to the fact that intersections between databases are not complete (i.e. some biological entities in database A may not exist in database B), in general terms the percentage of translated features has shown to be high and sufficient enough for pathway analysis purposes.

#### Linking metabolites and genes to KEGG identifiers – ‘Link Name to KEGGID’

In order to integrate transcriptomics and metabolomics data using common KEGG pathways, each omics data needs to be converted into the identifiers used by the KEGG database. Genes in KEGG are not directly attached to common identifiers such as RefSeq/PubChem IDs, rather, KEGG uses an independent identifier (a KEGG ID). Similarly, metabolite name conventions are diverse and KEGG uses their own compound IDs. The *Link Name to KEGGID* tool maps input gene/metabolite names to the KEGG database. The KEGG identifiers can then be retrieved and used to link to KEGG pathways. This process identifies which genes and metabolites are in shared pathways.

#### Features to Pathways – ‘Add KEGGID to Pathway Information’

Linking annotations form the data to a common identifier such as a KEGG identifier is the first step in an integrated the analysis. The next step is to link the KEGG identifiers to Pathways, which is a straight-forward parsing KEGG pathway files. When a feature maps to multiple pathways, multiple rows area created to indicate each relationship.

### Selection and filtering

We recommend performing a selection of metabolites/genes prior to the integration task. For example, differential expression can be used to filter metabolites/genes changing between treatments. Feature selection can be performed with any Galaxy tool that implements differential expression analysis. For example, for gene expression, the Galaxy implementations of edgeR^25^ or DEseq2^26^ can be used. For metabolomics, we recommend using SECIMTools^1^. These tools return lists of features (genes or metabolites) with associated p or q values that can be used for threshold-filtering.

### Integration Tool

#### Defining metabolite subsets

Metabolites can be analyzed as a function of gene expression as a whole data matrix, or as subsets of metabolites. Subsets can be created either by their annotation label or by their measured levels across samples. Annotation labels refer to the metabolite ID in the KEGG database, which can be assigned to multiple compounds of the input dataset. For example, when lipidomics data are processed, multiple compounds will be mapped to the KEGG ID sphingomyelin or ceramide, and hence these compounds are considered to be part of the same metabolite class. Metabolite groups can also be created based on their measurements across samples using a clustering technique. Here we use the SECIMTools^1^ implementation of MMC^21^ for unbiased clustering or metabolites. Note that subsets can be created by combining the annotation class with the MMC cluster or by a user defined knowledge base.

#### From genes in pathways to metagenes

Gene expression can be used a whole data matrix or genes can be selected that belong to specific pathways. An additional possibility is to concentrate pathway gene expression information in one or few metagenes that capture the variability pattern or “activity” of the pathway across samples. We have implemented the pathway metagene computation method described in^22^. This typically reduces the gene expression dataset to as many metagenes as annotated pathways.

#### Metabolite-Expression integration statistics

##### Unbiased analyses

To allow novel, unbiased discovery of gene-metabolite relationships, a simple correlation measure is implemented that calculates correlations between metabolite abundance and gene expression for all possible gene-metabolite pairs used as input. The tool then selects the top 500 correlation pairs to display data as an interaction network that can be further visually analyzed. Also the tool returns all correlation results in a tabular format for downstream analysis. This is a fully data-driven analysis that identifies the strongest co-variation relationships between genes and metabolites. Given the high number of correlations calculated here, it is important to estimate the potential for spurious association. GAIT does a simulation test, where the mean and variance of each compound/gene are assumed to be normal and this is used to generate observations at random for all features used. The simulated data are then processed identically to the observed data and the distribution of correlation coefficients are calculated. The simulation is performed 1000 times, and for each possible gene/metabolite pair, the frequency of random correlations above the correlation obtained with the original data is taken as p.value. Multiple testing may be adjusted for by several different already existing tools in Galaxy such as the ‘Multiple Testing Adjustment (MTA)’ tool implemented in SECIMTools^1^.

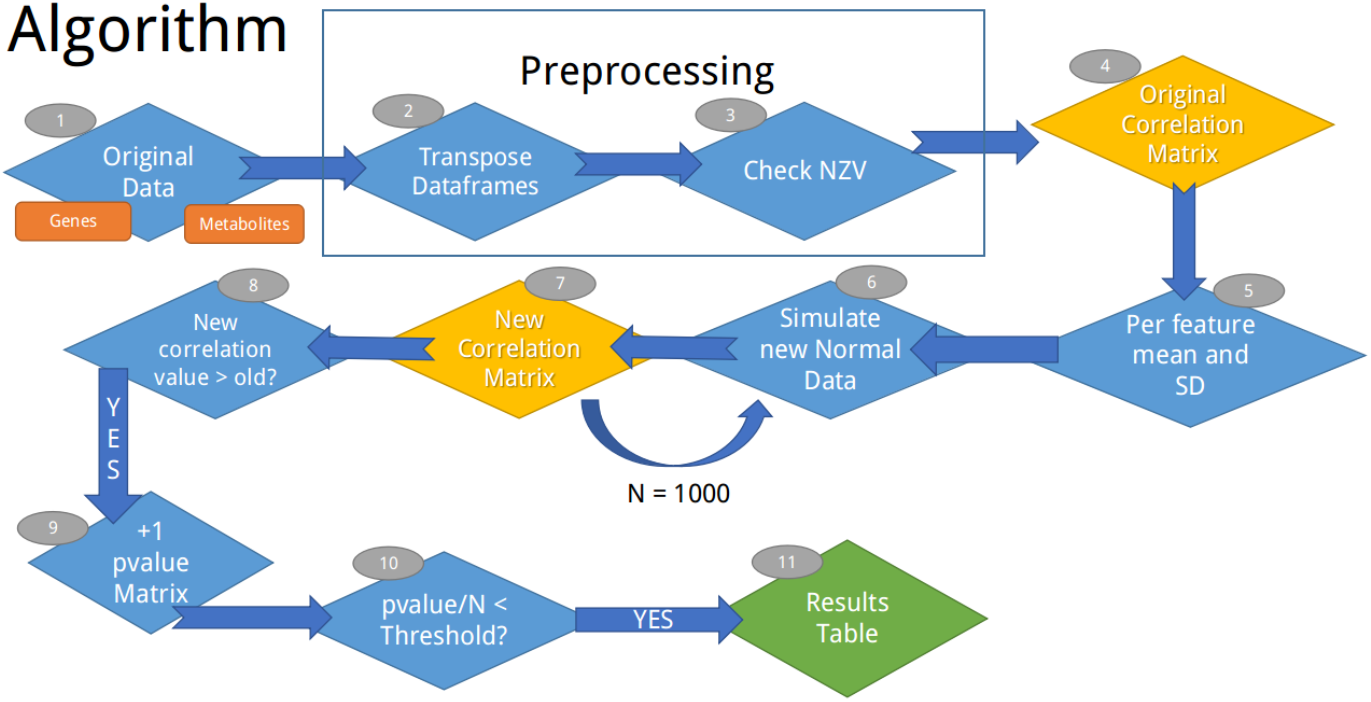

##### Direct Integration of metabolite quantity as a function of gene expression

We implement Sparse Partial Least Squares (sPLS) from the mixomics package^20^ as a method to explain metabolite level changes as a function of gene expression. In this approach, Gene Expression is the explanatory variable (X) and Metabolite levels is the response variable (Y). This statistical method can be applied to multiple combinations of Gene Expression and Metabolite matrices to create a highly flexible analysis framework where several regulation hypotheses can be tested.

###### a) Metabolite class vs. genes in associated pathways

A group of metabolites in the input data that map to the same compound ID in the KEGG database are considered a class and represent a single feature that might be present in one or several KEGG pathways. A pertinent question in this case is if all the metabolites in the same class are regulated in the same way or subsets of metabolites associate to different sets of genes. The sPLS model enables testing this hypothesis. By using as explanatory variables only genes in the pathway where the metabolite is annotated, the chance of identifying biologically interpretable associations is much higher.

###### c) Metabolite class vs. pathway metagenes

Annotation of metabolites to metabolic and signalling pathways is far from complete. In order to identify potential regulatory relationships beyond the pathways where metabolites are currently annotated a summary of expression in a given pathway can be derived as the combination of the expression profiles of genes in the pathway. These pathway metagenes can be fit as explanatory variables in sPLS models of groups of metabolites. When the response variables are metabolites in a class, this approach addresses the question of how the pathway activity network contributes to the regulation of the metabolites in the same class. When the response variables are co-expressed metabolites without a known annotation, associations with genes expressed in a pathway n may help identify the metabolites.

## Workflows

We have leveraged the power of the Galaxy platform, our new developed GAIT-TM tools and the existing contributions of the community to create complete Galaxy workflows for the integrated analysis of gene and metabolite expression data. These workflows are available on github (https://github.com/SECIMTools/gait-gm) and some examples are described here.

The ‘WF_gene_met_correlation’ Galaxy workflow implements the data-driven analysis described above. Stating with data files and feature identification information (e.g. m/z ratios or retention times), this workflow creates wide format datasets, design files, identifies the genes and metabolites of interest by ANOVA, annotates the genes and metabolites via KEGG and performs a correlation analysis between significant genes and metabolites to generate a table of correlation coefficients (Supplementary Figure 3). P-values for the correlation coefficients are calculated by simulating individual gene and metabolite datasets 1000 times using a normal distribution with means and standard deviations generated from the data. Sample size reflects the input datasets. Correlations are calculated on the simulated data. Correlations must be higher/lower than 95% of the randomly simulated values to be considered significant. An output image of this workflow is show in Supplementary Figure 3.

The ‘WF_int_met_class_2-genes_by_common_pathway’ Galaxy workflow implements one of the possible biologically-informed analyses described above. The workflow will create wide format datasets, design files, identify the genes and metabolites of interest by ANOVA, annotate the genes and metabolites via KEGG and integrate the gene expression and metabolite data by modelling metabolite classes as a function of the genes in the pathways where the metabolite is present. This approach is recommended as both gene expression and metabolite datasets are reduced to consider relationships that are likely to occur due to the pathway commonality of genes and metabolites. An output image of this workflow is show in Supplementary Figure 4.

Additional example Galaxy workflows are available (https://github.com/secimTools/gait-gm). For example, the ‘WF_int_met_2_metagene.ga’ workflow contains the same tools as described above but the options chosen in the ‘Metabolite – Gene Integration’ tool are different. In this case, the options select model metabolite classes as a function of metagenes that reflects the transcriptional activity of entire pathways. To include similarly behaving metabolites without regard to identification or type, the ‘WF_int_MMC_2_metagene.ga’ Galaxy workflow options implement the MMC tool to estimate modules that are modelled as a function of metagenes.

## Supporting information

Supplementary_Material_UserGuide

Supplementary_Material_Tool_Input_Output

## Authors’ contributions

LM conceived the research, supervised analyses and drafted the manuscript; FH programmed GAIT-GM tools, evaluated methods and performed data analysis; OM programmed GAIT-GM tools developed Conda recipes and tested all tools and submitted tools to pypi. MLl generated dataset used in the study, VF generated dataset used in the study, AMM, tested and debugged all tools, wrote all xml files, evaluated methods wrote documentation and the user guide and maintains the github. AC conceived the research, supervised analyses and drafted the manuscript. All authors read and approved the final manuscript.

## Acknowledgements

This work has been funded by National Institute of Health SECIM grant U24 DK097209 (LMM) and R03 CA222444 (AC, LMM). We are thankful of the support of HiPerGator and the University of Florida Research Computing Group.

