## Supplementary_Material_UserGuide for "GAIT-GM: Galaxy tools for modeling metabolite changes as a function of gene expression"

### GAIT-GM

**G**alaxy **A**nnotation and **I**ntegration **T**ools for **G**enes and **M**etabolites. These tools were designed to work with and complement the large set of Galaxy tools available for differential expression analysis and SECIMTools for metabolomics (https://github.com/secimTools/gait-gm) and therefore a complete workflow will require installing GAIT-GM, SECIMTools and the differential expression tools of choice. GAIT-GM tools can be run as python scripts (https://pypi.org/project/gait-gm/ ) or as part of a Galaxy installation.

#### Create: Design, Wide, and Annotation datasets from an Input wide dataset

This tool can be used to perform two tasks 1) convert a single file that contains data and annotation in wide format into two files in wide format, one with data and one with annotation 2) create a design file template that will be compatible with the wide data and annotation files. The tool will automatically check for a column containing unique feature identifiers (FeatureIDs). If no unique FeatureID column is located the tool will generate one. The user can specify a prefix for the unique FeatureID (e.g. 'met' for metabolite data). The tool creates a distinct Annotation Dataset containing the unique FeatureIDs (user-specified or generated by the tool) and any non-sample descriptor columns that were present in the input wide dataset (such as m/z ratio, retention time, compound name, etc.). The tool also creates a 'clean' Wide Dataset containing only samples in columns and features in rows. The Design Dataset is a template containing a single column called 'SampleID' with the names of the samples in the input Wide Dataset. This Design Dataset can be modified by the user to include metadata columns.

**Input:**

A TSV data file in wide format that may contain columns with annotation.

1. Select the **Input Dataset** from the drop-down menu.

2. If the **Input Dataset** has a column that specifies **unique FeatureIDs**, select **Yes**. If not, select **No**.

2a. If **Yes**, input the **name of the column in your Input Dataset that contains the unique FeatureIDs**.

2b. If **No**, choose a **prefix** (e.g. 'met'). All rows will be assigned a unique number after the prefix, separated by an underbar. If you chose not to use a prefix, the tool-created FeatureID column will be an underbar prepended to a unique number.

3. Select the **columns in your Input Dataset** that contain measured omics data. Columns that are not selected are treated as 'annotation' descriptor columns.

4. Click **Execute.**

**Output:**

Annotation Dataset TSV file: contains the **unique FeatureID column and any non-sample descriptor columns**.

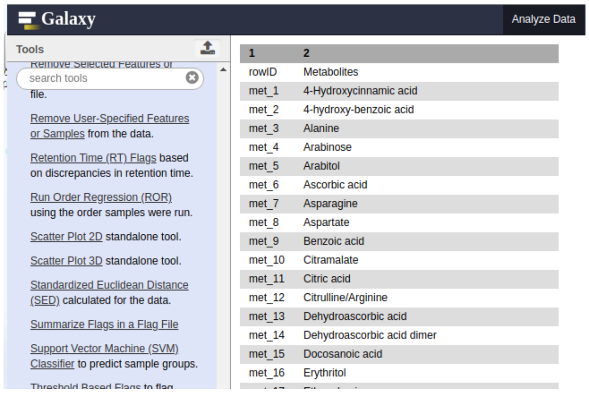


Design Dataset TSV template: contains a column called **sampleID** with the column headers from the input dataset selected in step 3. This file can be modified to include all necessary information for a design file.

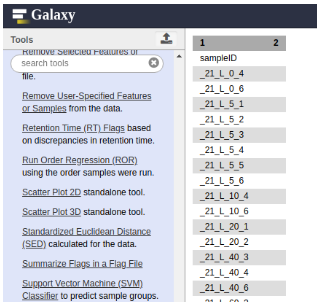


Wide Dataset: contains the **unique FeatureID column and all sample columns selected in Step 3**. Note that the output features line wrapping.

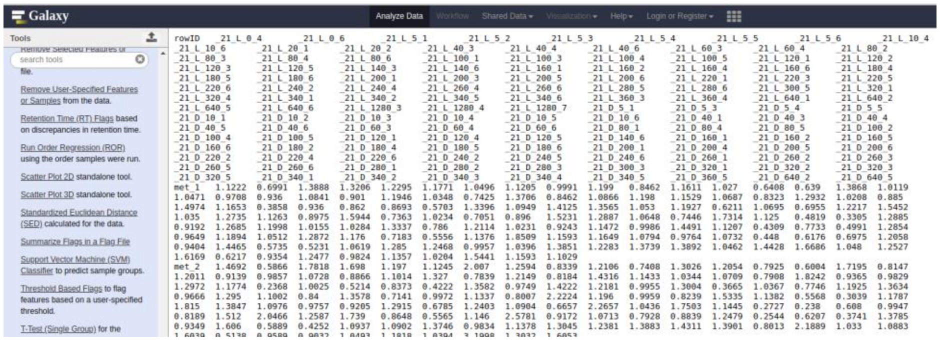


#### Map ENSEMBL IDs to Gene Symbols

This tool takes a dataset containing unique FeatureIDs and ENSEMBLIDs and provides gene symbols. The link from the ENSEMBLIDs to gene symbols is made using Biomart. The tool adds the following columns to the input dataset: GeneSymbol, Score, Selected, and Tie. The GeneSymbol column contains the short identifiers (typically 3 letter abbreviations) of the gene name. The Score column contains a value that represents how well the ENSEMBLID matched the returned GeneSymbol using the PyPi package Gene 3.0.0 (Wu, MacLeod and Su 2013). The Selected column = 'Yes' when an ENSEMBLID uniquely matches a GeneSymbol or when that row has the highest Score value. The Selected column = 'No' in the absence of a unique match for rows lower than the maximum score. If there is a tie in the Score the alphabetically first value is selected and the Tie column has a value of Yes. We note that FeatureID may not be unique in the resulting output dataset.

**Input:**

A TSV dataset containing a unique FeatureIDs and a column of ENSEMBLEIDs. Other columns may be present.

1. Select the **species** from which the ENSEMBLIDs are derived.

   Options include *Homo sapiens, Mus musculus, Rattus norvegicus, Drosophila melanogaster, Arabidopsis thaliana, Sacchoromyces cerevisiae, and Escherichia coli.*

2. Select the **dataset** containing the **FeatureID and associated ENSEMBLIDs**.

3. Enter the **name of the column** in your dataset that contains the **FeatureIDs**.

4. Enter the **name of the column** in your dataset that contains the **ENSEMBLEIDs**.

5. Click **Execute**.

**Output:**

A TSV dataset containing the data in the input dataset along with the following columns:

**GeneSymbol:** the gene symbol associated with a particular ENSEMBLEID. This is typically a 3 letter abbreviations of the gene name.

**Score:** a value that represents how well the ENSEMBLID matched the returned GeneSymbol.

**Selected:** Yes when ENSEMBLIDs uniquely match GeneSymbols, or when that row has the highest value of Score. Selected = No in the absence of a unique match for rows lower than the maximum score.

**Tie**: 'Yes' when FeatureIDs have matches with the same score.

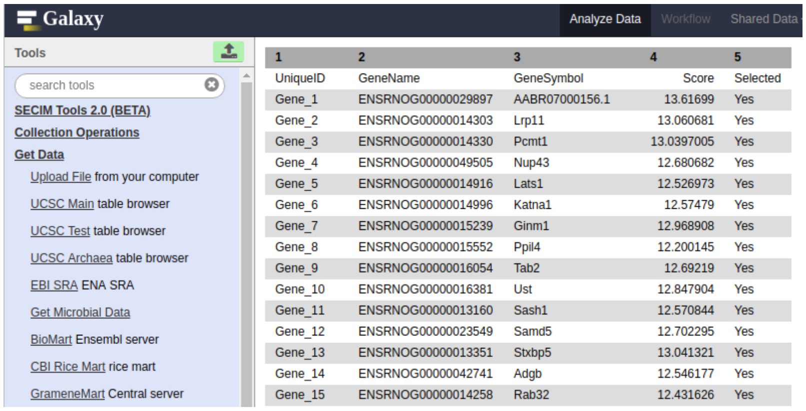


| 1 | 2 | 3 | 4 | 5 |
|---|---|---|---|---|
| UniqueID | GeneName | GeneSymbol | Score | Selected |
| Gene_1 | ENSRNOG00000029897 | AABR07000156.1 | 13.61699 | Yes |
| Gene_2 | ENSRNOG00000014303 | Lrp11 | 13.060681 | Yes |
| Gene_3 | ENSRNOG00000014330 | Pcmt1 | 13.0397005 | Yes |
| Gene_4 | ENSRNOG00000049505 | Nup43 | 12.680682 | Yes |
| Gene_5 | ENSRNOG00000014916 | Lats1 | 12.526973 | Yes |
| Gene_6 | ENSRNOG00000014996 | Katna1 | 12.57479 | Yes |
| Gene_7 | ENSRNOG00000015239 | Ginm1 | 12.968908 | Yes |
| Gene_8 | ENSRNOG00000015552 | Ppil4 | 12.200145 | Yes |
| Gene_9 | ENSRNOG00000016054 | Tab2 | 12.69219 | Yes |
| Gene_10 | ENSRNOG00000016381 | Ust | 12.847904 | Yes |
| Gene_11 | ENSRNOG00000013160 | Sash1 | 12.570844 | Yes |
| Gene_12 | ENSRNOG00000023549 | Samd5 | 12.702295 | Yes |
| Gene_13 | ENSRNOG00000013351 | Stxbp5 | 13.041321 | Yes |
| Gene_14 | ENSRNOG00000042741 | Adgb | 12.546177 | Yes |
| Gene_15 | ENSRNOG00000014258 | Rab32 | 12.431626 | Yes |

**Link compound and/or gene names in your Annotation Datast(s) to KEGG identifiers**

The 'Link Name to KEGGID' tool takes an annotation dataset containing metabolite compound names or gene symbols and links them to identifiers in KEGG (KEGGIDs) creating either a (a) Gene to KEGGID Link or a (b) Metabolite to KEGGID Link dataset. For gene expression data, the tool is designed to take the output from the 'Map ENSEMBLIDs to Gene Symbols' tool as input. If your input dataset contains a Selected column, the tool will link GeneSymbols to KEGGIDs where Selected = 'Yes'. Input Files without a Selected column must have a column containing unique FeatureIDs. This tool takes an annotation dataset containing unique FeatureIDs, ENSEMBLIDs (for gene expression data) and GeneSymbol/MetaboliteName and adds the following columns: 1) Name_in_KEGG, the name found in KEGG, 2) Matched, a column indicating whether a match was found in KEGG, 3) KEGGID, the KEGG identifier for the Match, 4) Score, a similarity score representing match similarity (calculated using the python SequenceMatcher class from difflib) and 5) a Tie column to indicate if a gene symbol or metabolite name matched more than one KEGGID.

User-specified metabolite names are linked to KEGGIDs by identifying the best match using the following procedure. Common metabolite prefixes are removed (cis-, trans-, d- , l- , (s)-, alpha-, beta-, alpha, beta, alpha-d-, beta-d-, alpha-l-, beta-l-, l-beta-, l-alpha-, d-beta-, d-alpha-). If the metabolite name given is an acid, then the name is modified to the conjugate base by replacing “ic acid”, “icacid” or “ic_acid” with “ate”. If amino acids are given in 1-letter or 3-letter abbreviations, names are modified to the full amino acid name. The following commonly used lipid abbreviations are modified to reflect the full names (SM = sphingomyelin, lysopc = lysophosphatidylcholine, PC = phosphatidylcholine, PE = phosphatidylethanolamine and LysoPE = lysophosphatidylethanolamine). Similarly, abbreviations for other commonly assayed metabolites are modified to reflect the full names (cit = citrate, orn = ornithine, thyr = thyroxine and boc = butoxycarbonyl). The code allows the addition of more synonyms. The user-specified metabolite names are retained in the output dataset for comparisons with the KEGG database.

Each parsed metabolite name is compared to metabolite names in KEGG. The best match in KEGG based on similarity score is returned. The similarity score (Score column) is based on the longest contiguous matching subsequence that does not contain 'junk' elements where 'junk' elements are defined as duplicates making up more than 1% of a sequence with minimum length of 200 (python SequenceMatcher class from difflib)

Selected = Yes for the match with the highest similarity score.

For metabolite names where the best match is tied with at least one other compound in KEGG, all matches are returned. A tie is determined as follows: if the Score is greater than 95% for 2 or more matches in the metabolite name then:

1) the Tie column = 'Yes' and a warning message will appear

2) the Selected column is sorted alphabetically on the Name_in_KEGG column. Note that the user-specified FeatureID and MetaboliteName may not be unique in the resulting output dataset.

**Input:**

Gene Annotation datasets must contain a column with gene symbols for matching in KEGG. Similarly,

Metabolomic Annotation datasets must contain a column with metabolite or compound names for matching in KEGG.

1. Select the **Species** from the drop-down menu.

Options include *Homo sapiens, Mus musculus, Rattus norvegicus, Drosophila melanogaster, Arabidopsis thaliana, Sacchoromyces cerevisiae, and Escherichia coli.*

2. If you have **both gene and metabolite Annotation datasets**, select 'Gene Expression + Metabolomic Annotation Datasets'.

-- Select your **Gene Expression Annotation dataset** from the drop-down menu.

-- Enter the name of the **column** in your Gene Expression Annotation dataset with the unique **FeatureIDs**.

-- Enter the name of the **column** in your Gene Expression Annotation dataset that contains the **Gene Symbols** to use for linking in KEGG.

-- Select your **Metabolite Annotation dataset** from the drop-down menu.

-- Enter the name of the **column** in your Metabolite Annotation dataset containing unique **FeatureIDs**.

-- Enter the name of the **column** in your Metabolite Annotation dataset that contains **the Metabolite names** to use for linking in KEGG.

**3. If you want to use a gene expression Annotation dataset only, select 'Gene Expression Annotation Dataset'.**

-- Select your **Gene Expression Annotation dataset** from the drop-down menu.

-- Enter the name of the **column** in your Gene Expression Annotation dataset containing unique **FeatureIDs**.

-- Enter the name of the **column** in your Gene Expression Annotation dataset that contains the **Gene Symbols** to use for linking in KEGG.

**4. If you want to use a metabolite Annotation dataset only, select 'Metabolomic Annotation Dataset'.**

-- Select your **Metabolite Annotation dataset** from the drop-down menu.

-- Enter the name of the **column** in your Metabolite Annotation dataset containing unique **FeatureIDs**.

-- Enter the name of the **column** in your Metabolite Annotation dataset that contains the compound names to use for linking in KEGG.

5. Click **Execute**.

**Output:**

For Gene Expression or Metabolite datasets the output (a **Gene to KEGGID Link dataset** or **Metabolite to KEGGID Link dataset**) consists of a TSV dataset containing the information from the original input file with the following columns added:

**Feature_Type:** column indicating whether matching was carried out for metabolites or genes.

**Matched:** column indicating whether a match in KEGG was found. Yes/No

**Name_in_KEGG**: column containing the KEGG name for the match.

**KEGGID:** column containing the KEGG identifier for the match.

**Similarity:** value indicating the similarity between the user-specified feature and the match in KEGG. Ranges from 0 to 1. Similarity is based on the longest contiguous matching subsequence that does not contain 'junk' elements where 'junk' elements are defined as duplicates making up more than 1% of a sequence with minimum length of 200 (python SequenceMatcher class from difflib (https://docs.python.org/2/_sources/library/difflib.rst.txt, module and section author Tim Peters)).

**Tie:** In cases where multiple matches are found for a given featureID, Tie = Yes if the similarity is greater than 99%.

**Selected**: For features with multiple matches and different similarity scores, the Selected column = Yes for the match with the highest similarity score. For features with multiple matches and the same similarity score, the Selected column = Yes for the first compound in an alphabetized list.

An example of a Gene to KEGGID Link output dataset:

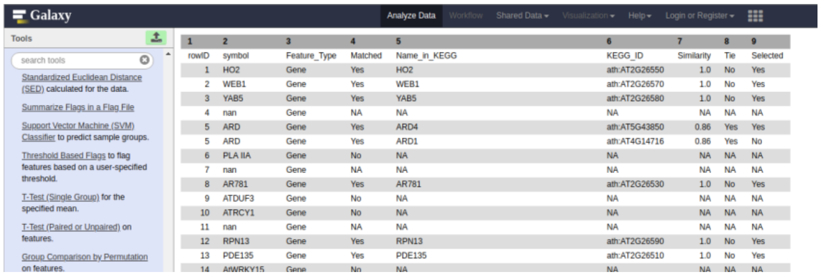

| 1 | 2 | 3 | 4 | 5 | 6 | 7 | 8 | 9 |
|---|---|---|---|---|---|---|---|---|
| rowID | symbol | Feature_Type | Matched | Name_in_KEGG | KEGG_ID | Similarity | Tie | Selected |
| 1 | HO2 | Gene | Yes | HO2 | ath:AT2G26550 | 1.0 | No | Yes |
| 2 | WEB1 | Gene | Yes | WEB1 | ath:AT2G26570 | 1.0 | No | Yes |
| 3 | YAB5 | Gene | Yes | YAB5 | ath:AT2G26580 | 1.0 | No | Yes |
| 4 | nan | Gene | NA | NA | NA | NA | NA | NA |
| 5 | ARD | Gene | Yes | ARD4 | ath:AT5G43850 | 0.86 | Yes | Yes |
| 5 | ARD | Gene | Yes | ARD1 | ath:AT4G14716 | 0.86 | Yes | No |
| 6 | PLA IIA | Gene | No | NA | NA | NA | NA | NA |
| 7 | nan | Gene | NA | NA | NA | NA | NA | NA |
| 8 | AR781 | Gene | Yes | AR781 | ath:AT2G26530 | 1.0 | No | Yes |
| 9 | ATDUF3 | Gene | No | NA | NA | NA | NA | NA |
| 10 | ATRCY1 | Gene | No | NA | NA | NA | NA | NA |
| 11 | nan | Gene | NA | NA | NA | NA | NA | NA |
| 12 | RPN13 | Gene | Yes | RPN13 | ath:AT2G26590 | 1.0 | No | Yes |
| 13 | PDE135 | Gene | Yes | PDE135 | ath:AT2G26510 | 1.0 | No | Yes |
| 14 | AtWRKY15 | Gene | No | NA | NA | NA | NA | NA |

An example of a Metabolite to KEGGID Link output dataset:

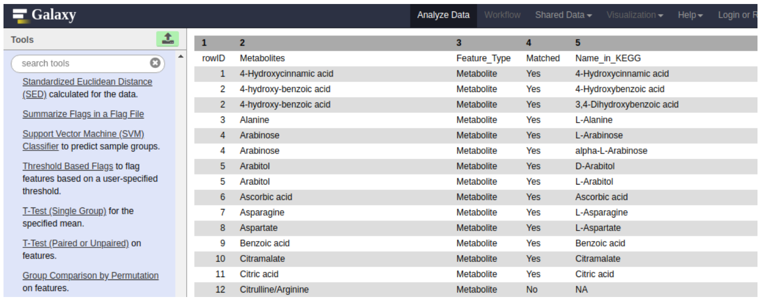

| 1 | 2 | 3 | 4 | 5 |
|---|---|---|---|---|
| rowID | Metabolites | Feature_Type | Matched | Name_in_KEGG |
| 1 | 4-Hydroxycinnamic acid | Metabolite | Yes | 4-Hydroxycinnamic acid |
| 2 | 4-hydroxy-benzoic acid | Metabolite | Yes | 4-Hydroxybenzoic acid |
| 2 | 4-hydroxy-benzoic acid | Metabolite | Yes | 3,4-Dihydroxybenzoic acid |
| 3 | Alanine | Metabolite | Yes | L-Alanine |
| 4 | Arabinose | Metabolite | Yes | L-Arabinose |
| 4 | Arabinose | Metabolite | Yes | alpha-L-Arabinose |
| 5 | Arabitol | Metabolite | Yes | D-Arabitol |
| 5 | Arabitol | Metabolite | Yes | L-Arabitol |
| 6 | Ascorbic acid | Metabolite | Yes | Ascorbic acid |
| 7 | Asparagine | Metabolite | Yes | L-Asparagine |
| 8 | Aspartate | Metabolite | Yes | L-Aspartate |
| 9 | Benzoic acid | Metabolite | Yes | Benzoic acid |
| 10 | Citramalate | Metabolite | Yes | Citramalate |
| 11 | Citric acid | Metabolite | Yes | Citric acid |
| 12 | Citrulline/Arginine | Metabolite | No | NA |

**Add KEGG Pathway Information using KEGGIDs.**

The 'Add KEGG Pathway Information' tool takes a Gene to KEGG Link dataset, a Metabolomic to KEGG Link dataset or both and adds KEGG Pathway Names using KEGGIDs. The tool was designed to take the output from the "Link Name to KEGGID' tool as input (for example the Gene to KEGGID Link dataset) but other datasets containing KEGGIDs can be used as well.

The user will get different outputs from the 'Add KEGG Pathway Info Tool', depending on the input. If a Gene to KEGGID Link dataset is given as input the tool outputs the following three files: 1) a Gene KEGG Pathway dataset containing the FeatureID, Feature_Name, Feature_Type and KEGGID columns from the input file and KEGG_PathwayIDs and KEGG Pathway Names from KEGG, 2) a GeneKeggID2PathwayID dataset containing all gene KEGGIDs in KEGG and their associated pathway KEGGIDs and 3) a PathwayID2PathwayNames dataset containing all of the pathway KEGGIDs and their associated KEGG pathway names.

Analoguous files are generated by the tool if a Metabolite to KEGGID Link dataset is input by the user.

Note: FeatureIDs and KEGGIDs may not be unique in the output.

1. Select your **Species** from the drop-down menu.

Options include *Homo sapiens*, *Mus musculus*, *Rattus norvegicus*, *Drosophila melanogaster, Arabidopsis thaliana, Sacchoromyces cerevisiae, and Escherichia coli.*

2a. If you have both Gene Expression and Metabolite Annotation datasets, **select 'Gene Expression + Metabolomic Annotation datasets'**.

-- Select your **Gene to KEGGID Link dataset** from the drop-down menu.

-- Enter the name of the **column** in your Gene to KEGGID Link dataset with the unique **geneIDs**.

-- Enter the name of the **column** in your Gene to KEGGID Link dataset that contains **gene symbols**.

-- Enter the name of the **column** in your Gene to KEGGID Link dataset that contains the **KEGG identifiers**.

-- Select **your Metabolite to KEGGID Link dataset** from the drop-down menu.

-- Enter the name of the **column** in your Metabolite to KEGGID Link dataset with the unique **FeatureIDs**.

-- Enter the name of the **column** in your Metabolite to KEGGID Link dataset that contains **metabolite names**.

-- Enter the name of the **column** in your Metabolite to KEGGID Link dataset that contains the **KEGG identifiers**.

3a. Click **Execute**.

2b. If you want to use a Gene Expression Annotation File only, select **'Gene Expression Annotation dataset'**.

-- Select **your Gene to KEGGID Link dataset** from the drop-down menu.

-- Enter the name of the **column** in your Gene to KEGGID Link dataset with the unique **geneIDs**.

-- Enter the name of the **column** in your Gene to KEGGID Link dataset that contains **gene symbols**.

-- Enter the name of the **column** in your Gene to KEGGID Link dataset File that contains the **KEGG identifiers**.

3b. Click **Execute**.

2c. If you want to use a Metabolite Annotation File only, select **'Metabolomic Annotation Dataset'**.

-- Select your **Metabolite to KEGGID Link dataset** from the drop-down menu.

-- Enter the name of the **column** in your Metabolite to KEGGID Link dataset with the unique **FeatureIDs**.

-- Enter the name of the **column** in your Metabolite to KEGGID Link dataset that contains **metabolite names**.

-- Enter the name of the **column** in your Metabolite to KEGGID Link dataset that contains the **KEGG identifiers**.

**3c. Click Execute.**

**Input:**

The Gene to KEGGID Link and/or Metabolite to KEGGID Link TSV Files must contain the following 3 columns;

1) **unique FeatureIDs**, 2) **gene or metabolite names** and 3) **KEGGIDs**. Other columns may be present. Note that input files do not have to be generated by the 'Link Name to KEGGID' tool.

**Output:**

The user will get different output from the 'Add KEGG Pathway Information' tool, depending on whether they include the 'Gene to KEGGID Link' File, the 'Metabolite to KEGGID Link' File, or both.

The following files are generated if the **'Gene to KEGGID Link**' File is selected:

**1. Gene KEGG Pathway.** TSV file containing the **FeatureID**, **GeneName**, **Feature_Type** and **KEGGID** columns from the input dataset along with the following additional columns:

**PathwayID**: the KEGG Pathway identifiers for the KEGGIDs

**Pathway_Name**: the KEGG Pathway name for the KEGGIDs.

Analyze Data Workflow Shared Data Visualization Help Login or Register

| 1 | 2 | 3 | 4 | 5 | 6 |
|---|---|---|---|---|---|
| UniqueID | Feature_Name | Feature_Type | KEGG_ID | Pathway_ID | Pathway_Name |
| 1 | HO2 | Gene | ath:AT2G26550 | NA | NA |
| 2 | WEB1 | Gene | ath:AT2G26570 | NA | NA |
| 3 | YAB5 | Gene | ath:AT2G26580 | NA | NA |
| 4 | nan | Gene | NA | NA | NA |
| 5 | ARD | Gene | ath:AT5G43850 | path:ath00270 | Cysteine and methionine metabolism |
| 5 | ARD | Gene | ath:AT5G43850 | path:ath01100 | Metabolic pathways |
| 6 | PLA IIA | Gene | NA | NA | NA |
| 7 | nan | Gene | NA | NA | NA |
| 8 | AR781 | Gene | ath:AT2G26530 | NA | NA |
| 9 | ATDUF3 | Gene | NA | NA | NA |
| 10 | ATRCY1 | Gene | NA | NA | NA |
| 11 | nan | Gene | NA | NA | NA |
| 12 | RPN13 | Gene | ath:AT2G26590 | path:ath03050 | Proteasome |
| 13 | PDE135 | Gene | ath:AT2G26510 | NA | NA |
| 14 | AtWRKY15 | Gene | NA | NA | NA |
| 15 | nan | Gene | NA | NA | NA |
| 16 | SMU2 | Gene | ath:AT2G26460 | NA | NA |
| 17 | nan | Gene | NA | NA | NA |
| 18 | UGT76D1 | Gene | ath:AT2G26480 | NA | NA |
| 19 | RHC2A | Gene | ath:AT2G39720 | NA | NA |
| 20 | CYT1 | Gene | ath:AT2G39770 | path:ath00051 | Fructose and mannose metabolism |
| 20 | CYT1 | Gene | ath:AT2G39770 | path:ath00520 | Amino sugar and nucleotide sugar metabolism |
| 20 | CYT1 | Gene | ath:AT2G39770 | path:ath01100 | Metabolic pathways |
| 20 | CYT1 | Gene | ath:AT2G39770 | path:ath01110 | Biosynthesis of secondary metabolites |
| 21 | RCA | Gene | ath:AT2G39730 | NA | NA |
| 22 | ATBPM3 | Gene | NA | NA | NA |

2. **GeneKeggID2PathwayID.** A downloaded file from KEGG that contains **ALL** the **gene KEGGIDs** and associated pathway identifiers (**PathwayIDs**) for the selected species.

| 1 | 2 |
|---|---|
| path:ath00010 | ath:AT1G01090 |
| path:ath00010 | ath:AT1G09780 |
| path:ath00010 | ath:AT1G09870 |
| path:ath00010 | ath:AT1G12000 |
| path:ath00010 | ath:AT1G13440 |
| path:ath00010 | ath:AT1G16300 |
| path:ath00010 | ath:AT1G20950 |
| path:ath00010 | ath:AT1G22170 |
| path:ath00010 | ath:AT1G22430 |
| path:ath00010 | ath:AT1G22440 |
| path:ath00010 | ath:AT1G23190 |
| path:ath00010 | ath:AT1G23800 |
| path:ath00010 | ath:AT1G24180 |
| path:ath00010 | ath:AT1G30120 |
| path:ath00010 | ath:AT1G32440 |

3. **PathwayID2pathwayNames.** A downloaded file from KEGG that contains **ALL** the KEGG pathway identifiers (**PathwayID**s) and the associated KEGG pathway names (**Pathway_Name**) for the selected species.

| 1 | 2 |
|---|---|
| path:ath00010 | Glycolysis / Gluconeogenesis - Arabidopsis thaliana (thale cress) |
| path:ath00020 | Citrate cycle (TCA cycle) - Arabidopsis thaliana (thale cress) |
| path:ath00030 | Pentose phosphate pathway - Arabidopsis thaliana (thale cress) |
| path:ath00040 | Pentose and glucuronate interconversions - Arabidopsis thaliana (thale cress) |
| path:ath00051 | Fructose and mannose metabolism - Arabidopsis thaliana (thale cress) |
| path:ath00052 | Galactose metabolism - Arabidopsis thaliana (thale cress) |
| path:ath00053 | Ascorbate and aldarate metabolism - Arabidopsis thaliana (thale cress) |
| path:ath00061 | Fatty acid biosynthesis - Arabidopsis thaliana (thale cress) |
| path:ath00062 | Fatty acid elongation - Arabidopsis thaliana (thale cress) |
| path:ath00071 | Fatty acid degradation - Arabidopsis thaliana (thale cress) |
| path:ath00072 | Synthesis and degradation of ketone bodies - Arabidopsis thaliana (thale cress) |
| path:ath00073 | Cutin, suberine and wax biosynthesis - Arabidopsis thaliana (thale cress) |
| path:ath00100 | Steroid biosynthesis - Arabidopsis thaliana (thale cress) |
| path:ath00130 | Ubiquinone and other terpenoid-quinone biosynthesis - Arabidopsis thaliana (thale cress) |
| path:ath00190 | Oxidative phosphorylation - Arabidopsis thaliana (thale cress) |
| path:ath00195 | Photosynthesis - Arabidopsis thaliana (thale cress) |

The following files are generated if the '**Metabolite to KEGGID Link**' File is selected:

**1. Metabolite KEGG Pathway File**. TSV file containing the **FeatureID**, **metabolite name**, **Feature_Type** and **KEGGID** columns from the input dataset along with the following additional columns:

**PathwayID**: the KEGG Pathway identifiers for the KEGGIDs

**Pathway_Name**: the KEGG Pathway names for the KEGGIDs.

2. **MetaboliteKeggID2PathwayID**. Downloaded file from KEGG that contains **ALL** the **metabolite KEGGIDs** and associated pathway identifiers (**PathwayIDs**) for the selected species.

3. **PathwayID2pathwayNames.** A downloaded file from KEGG that contains **ALL** the KEGG pathway identifiers (**PathwayID**s) and the associated KEGG pathway names (**Pathway_Name**) for the selected species.

If '**Gene Expression + Metabolomic Datasets**' is selected, then output for both gene expression and metabolites is generated by the tool. Note that since the PathwayID2pathwayNames file is the same for both gene expression and metabolite datasets, only one file is generated.

#### <u>Add Binary (0/1) P-value Flags</u>

This tool generates an indicator variable (0 or 1) to identify p-values below a user-specified threshold. A "1" is used to indicate (flag) p-values less than the indicated threshold p-value. The user can flag nominal p-values or p-values after correction for multiple testing.

**Input:**

A TSV dataset containing P-values in a single column.

1. Select the **Dataset containing the P-values** you wish to flag from your History.

2. Enter the name of the **column** in your Dataset that contains the unique **FeatureIDs**.

3. Enter the name of the **column** in your Dataset that contains the **p-values**.

3. Enter the **p-value threshold(s)** for flagging. P-values less than the given threshold(s) will be flagged with a 1. If you enter more than 1 threshold value, separate the values with a comma (no spaces). Default values are 0.1, 0.05, and 0.01.

**Output:**

**1. Output File.** A TSV file containing **the same columns as the Input Dataset plus** additional column(s) containing **0/1 binary indicators** for whether the p-value was less that the user-specified threshold. The indicator columns are named by appending the user-specified threshold to 'Flag_' prefix (e.g. Flag_*user-specified threshold*, Flag_0.10*)*.

Analyze Data | Workflow | Shared Data | Visualization | Help | Login or Register

| 1 | 2 | 3 | 4 | 5 | 6 | 7 | 8 | 9 |
|---|---|---|---|---|---|---|---|---|
| UniqueID | MetName | logFC | AveExpr | t | P.Value | adj.P.Val | B | Flag_0.15 |
| Met_6 | SM C16:1 | 0.184563022813792 | 12.1314574685901 | 3.19422607775338 | 0.0099941857347226 | 0.456883398574827 | -2.53695037527621 | 1 |
| Met_126 | Spermidine | -0.0691475900899725 | 13.1683074598369 | -3.03265871016923 | 0.0130969384852503 | 0.456883398574827 | -2.79485549278087 | 1 |
| Met_107 | DOPA | -0.181632471939677 | 13.3178495402036 | -2.92528956820157 | 0.015688969318725 | 0.456883398574827 | -2.90554528935395 | 1 |
| Met_36 | PC aa C34:4 | 0.132428542778042 | 12.2180308925578 | 2.78205014553766 | 0.0199787307844689 | 0.456883398574827 | -3.00789053463321 | 1 |
| Met_111 | Gly | -0.118479527476564 | 13.9404685051305 | -2.66164293130856 | 0.0244901253745921 | 0.456883398574827 | -3.16711249695809 | 1 |
| Met_117 | Lys | -0.12725044551123 | 13.4132926037154 | -2.56066114614138 | 0.0290522589565527 | 0.456883398574827 | -3.42178691022983 | 1 |
| Met_42 | PC aa C36:5 | 0.073817416404971 | 12.7118154583614 | 2.55769269736229 | 0.0291984831282408 | 0.456883398574827 | -3.51628265235612 | 1 |
| Met_8 | SM C18:1 | 0.164977077093731 | 13.0694110587293 | 2.39165515909878 | 0.0386517851077047 | 0.456883398574827 | -3.81184713722168 | 1 |
| Met_75 | PC ae C36:5 | 0.0369147179734277 | 12.8042445927054 | 2.38550149976137 | 0.0386733515074654 | 0.456883398574827 | -3.79747639668756 | 1 |
| Met_113 | Histamine | -0.259678852093632 | 13.0358215747059 | -2.25130374235808 | 0.0489445843965551 | 0.456883398574827 | -4.0188604638993 | 1 |
| Met_48 | PC aa C38:5 | 0.0418402628631966 | 13.3090425242909 | 2.23449775706251 | 0.0499258731164521 | 0.456883398574827 | -3.96631013903615 | 1 |
| Met_100 | ADMA | -0.195179955519993 | 13.1701218869216 | -2.23863383733283 | 0.0499955518089795 | 0.456883398574827 | -4.01906447714316 | 1 |
| Met_73 | PC ae C36:3 | -0.0276789949809828 | 12.9734932278677 | -2.13431109674463 | 0.0590807019727391 | 0.456883398574827 | -4.2114499142652 | 1 |
| Met_30 | PC aa C32:1 | 0.0433713013174106 | 13.3200094173184 | 2.11275238940995 | 0.0612517217367824 | 0.456883398574827 | -4.14104358700268 | 1 |

**2. Flags File.** A TSV file containing only the **FeatureID** column from the input dataset and the tool generated **binary indicator flags**.

| 1 | 2 |
|---|---|
| UniqueID | Flag_0.15 |
| Met_6 | 1 |
| Met_126 | 1 |
| Met_107 | 1 |
| Met_36 | 1 |
| Met_111 | 1 |
| Met_117 | 1 |
| Met_42 | 1 |
| Met_8 | 1 |
| Met_75 | 1 |
| Met_113 | 1 |
| Met_48 | 1 |

**Metabolite – Gene Correlation**

The tool performs a correlation analysis between genes (in a gene expression wide dataset) and metabolites (in a metabolite wide dataset) to generate a table of correlation coefficients. P-values for the correlation coefficients are calculated by simulating individual gene and metabolite datasets 1000 times using a normal distribution with means and standard deviations generated from the data. Sample size reflects the input datasets. Correlations are calculated on the simulated data. Correlations must be higher/lower than 95% of the randomly simulated values to be considered significant.

**Input:**

The tool requires 2 input datasets, 1) a Gene Expression Wide Dataset containing gene expression measurements for each sample and 2) a Metabolite Wide Dataset containing metabolite measurements for the same samples. Wide datasets, containing a unique FeatureID column and samples as additional columns, can be generated using the 'Create: Design, Wide and Annotation' tool. Sample names are user-specified but the sample names in the two files must match. Optional inputs include a Gene and or Metabolite Annotation File(s) (TSV format) containing a column for the FeatureID (genes or metabolites) and a second column with the desired feature label for the graphical output.

1. Select **the Gene Expression Wide Dataset** from your History.

2. Enter the name of the **column** in your Gene Expression Wide Dataset that contains the unique **FeatureIDs**.

3. Select whether you want to use **a Gene Expression Annotation File**. If 'Yes' then the user can chose a column in an Annotation File to use for labeling output files so they are more user-friendly.

– if you select **'Yes'**

– Select the **Gene Expression Annotation File** from your History.

– Enter the name of the **column** in your Gene Expression Annotation File that you want to use to **label** the output files.

4. Select the **Metabolite Wide Dataset** from your History.

5. Enter the name of the **column** in your Metabolite Wide Dataset that contains the unique **FeatureIDs**.

6. Select whether you want to use **a Metabolite Annotation File**. If 'Yes' then the user can chose a column in an Annotation File to use for labeling output files so they are more user-friendly.

– if you select **'Yes'**

– Select **the Metabolite Annotation File** from your History.

– Enter the name of the **column** in your Metabolite Annotation File that you want to use to **label** the output files.

7. Select a **Correlation method**. Options include Pearson (Pearson’s standard correlation coefficient, default value), Spearman (Spearman’s rank correlation) or Kendall (Kendall’s Tau correlation coefficient).

8. Enter a **p-value threshold**. This value is used to filter the results in the 'Correlation File' to only those correlations whose P-value is less than this user-specified threshold. Entering a P-value threshold of 1 will output all correlation coefficients.

9. Click **Execute**.

**Output:**

The tool outputs 2 TSV files and a PDF figure.

1. The **Correlation File**. Contains gene-metabolite correlations with p-values less than the user-specified threshold. Note that a threshold of 1 outputs all coefficients. The Correlation File is sorted by the absolute value of the correlation coefficient.

Analyze Data Workflow Shared Data Visualizati

| 1 | 2 | 3 | 4 |
|---|---|---|---|
| "gene" | "metabolite" | "correlation" | "p-value" |
| Gene_3309: ENSRNOG00000049378 | Met_8: SM C18:1 | 0.983849911407111 | 0 |
| Gene_3309: ENSRNOG00000049378 | Met_28: PC aa C30:0 | 0.982989437292042 | 0 |
| Gene_6670: ENSRNOG00000004936 | Met_65: PC ae C32:2 | -0.982917671463121 | 0 |
| Gene_8206: ENSRNOG00000032659 | Met_61: PC ae C30:0 | 0.980127890041639 | 0 |
| Gene_7256: ENSRNOG00000026589 | Met_61: PC ae C30:0 | 0.979163093390531 | 0 |
| Gene_3511: ENSRNOG00000005668 | Met_64: PC ae C32:1 | 0.974678661238214 | 0 |
| Gene_3987: ENSRNOG00000004810 | Met_119: NTyr | 0.973603247367976 | 0 |
| Gene_10978: ENSRNOG00000016123 | Met_117: Lys | -0.972844557406263 | 0 |
| Gene_664: ENSRNOG00000031662 | Met_61: PC ae C30:0 | 0.972317780500125 | 0 |
| Gene_3892: ENSRNOG00000000172 | Met_65: PC ae C32:2 | -0.969636290757066 | 0 |
| Gene_2779: ENSRNOG00000012826 | Met_61: PC ae C30:0 | 0.969425648757226 | 0 |
| Gene_13156: ENSRNOG00000056040 | Met_61: PC ae C30:0 | -0.968384709823354 | 0 |
| Gene_920: ENSRNOG00000012206 | Met_61: PC ae C30:0 | 0.9674572532453 | 0 |
| Gene_4482: ENSRNOG00000039530 | Met_61: PC ae C30:0 | -0.967240371229274 | 0 |
| Gene_1571: ENSRNOG00000016883 | Met_14: C16:1 | 0.966334736501741 | 0 |
| Gene_8860: ENSRNOG00000007125 | Met_61: PC ae C30:0 | -0.965920445960242 | 0 |
| Gene_13146: ENSRNOG00000025998 | Met_53: PC aa C40:4 | 0.965860602911394 | 0 |
| Gene_58: ENSRNOG00000011720 | Met_12: SM C26:1 | 0.965470421009984 | 0 |
| Gene_9837: ENSRNOG00000007869 | Met_61: PC ae C30:0 | 0.963904874852478 | 0 |
| Gene_6161: ENSRNOG00000005292 | Met_117: Lys | -0.963580445123533 | 0 |

2. **Correlation Matrix**. The correlation results in matrix format.

Analyze Data Workflow Shared Data Visualization Help Login or Register

This dataset is large and only the first megabyte is shown below.
Show all | Save

| | Met_1: SM (OH) C16:1 | Met_2: SM (OH) C22:1 | Met_3: SM (OH) C22:2 | Met_4: SM (OH) C24:1 | Met_5: SM C16:0 Met_6: SM C16:1 | Met_7: SM C18: |
|---|---|---|---|---|---|---|
| Gene_1: | ENSRNOG00000029897 | 0.159010583779719 | 0.12963093780143 | -0.153417845171639 | -0.157661743754091 | -0.0236477644671398 |
| Gene_2: | ENSRNOG00000014303 | 0.176947560106124 | 0.760228061514014 | 0.504385873070097 | 0.688285258409744 | 0.0620797449255194 |
| Gene_3: | ENSRNOG00000014330 | 0.569726435947249 | -0.211085485804449 | -0.409938791132407 | -0.264297285676802 | 0.430713233617065 |
| Gene_4: | ENSRNOG00000049505 | -0.206947863930029 | -0.797892834624819 | -0.713612458990072 | -0.898697557628883 | 0.188293344973583 |
| Gene_5: | ENSRNOG00000014916 | -0.291696727378155 | 0.00915566570080262 | 0.363788808858826 | 0.128180335373216 | 0.111384263441703 |
| Gene_6: | ENSRNOG00000014996 | -0.394801108792586 | -0.182611725718792 | 0.155719239415693 | 0.157892870888078 | -0.0986350007448756 |
| Gene_7: | ENSRNOG00000015239 | -0.236207501568188 | -0.355927511880879 | -0.207967462080168 | -0.546247461084066 | 0.274843995361284 |
| Gene_8: | ENSRNOG00000015552 | -0.693631454725884 | -0.114718611460565 | 0.331445206465841 | 0.236643156246417 | -0.689791120081257 |
| Gene_9: | ENSRNOG00000016054 | 0.141834019434679 | -0.125459827241421 | -0.329676362393142 | -0.18765900017901 | -0.128124661236994 |
| Gene_10: | ENSRNOG00000016381 | 0.0940126145280227 | -0.743042089580945 | -0.777007156198828 | -0.780893346451167 | 0.463753054302198 |
| Gene_11: | ENSRNOG00000013160 | -0.000730789684253313 | -0.180692325642632 | -0.171857426512291 | -0.162525406865839 | 0.0083032399458998 |
| Gene_12: | ENSRNOG00000023549 | -0.0291841311328149 | 0.066606778277722 | -0.235211837955011 | -0.208607102891644 | -0.209936561382832 |
| Gene_13: | ENSRNOG00000013351 | -0.0754157454308071 | -0.00910064107744299 | 0.242767478799907 | 0.321934529719947 | 0.128165851658872 |
| Gene_14: | ENSRNOG00000042741 | 0.0272575745604586 | 0.391132129960196 | 0.159117595221318 | 0.337059235357512 | 0.217286712357083 |
| Gene_15: | ENSRNOG00000014258 | 0.281968198827613 | -0.168136411800489 | -0.375904015192005 | -0.216589869690646 | 0.405726011873312 |
| Gene_16: | ENSRNOG00000014290 | -0.190439284399228 | -0.149555638156026 | 0.202990039452886 | 0.184767109802283 | 0.0549433530864779 |
| Gene_17: | ENSRNOG00000014450 | -0.171072821300361 | 0.850426945304106 | 0.638588383669237 | 0.61172434633455 | -0.398136127412687 |
| Gene_18: | ENSRNOG00000014852 | -0.483216611014956 | 0.0981484341764666 | 0.221460516980381 | -0.0317994853966336 | -0.429645967164994 |
| Gene_19: | ENSRNOG00000040242 | 0.187558033158953 | -0.872524535198081 | -0.879470210199532 | -0.737857430109942 | 0.115673555403745 |
| Gene_20: | ENSRNOG00000011058 | 0.57641353334891 | 0.470643739318964 | 0.0616240050612764 | 0.0911267571999317 | 0.454661334511674 |
| Gene_21: | ENSRNOG00000014908 | -0.261721238539743 | 0.0915656978984286 | 0.19254516280949 | 0.193408633520277 | -0.40724145969753 |
| Gene_22: | ENSRNOG00000025587 | -0.235518817278243 | -0.127988640189381 | -0.113689268061362 | -0.292617419745446 | 0.0811736884857827 |
| Gene_23: | ENSRNOG00000015217 | 0.0678312341257502 | -0.131357514968195 | -0.163486048020789 | -0.214110598062642 | 0.501174487155058 |
| Gene_24: | ENSRNOG00000015473 | 0.40524187075917 | -0.279196780391663 | -0.193007082965765 | -0.0980555085800478 | 0.335663102120411 |
| Gene_25: | ENSRNOG00000015551 | -0.0429363443547737 | 0.114025275888278 | -0.055297499562428 | -0.128364290459116 | -0.486915436440036 |

3. **Correlation Figure**. A network representation of the top 500 gene-metabolite correlations based on the absolute value of the correlation coefficient. Max number of correlation coefficients is 500.

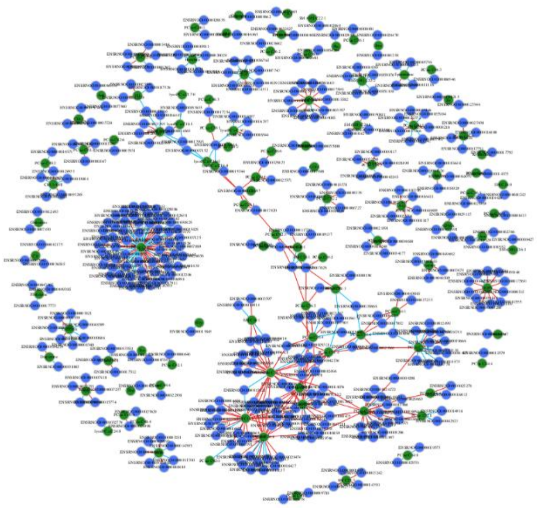

#### Metabolite – Gene integration

This tool carries out the integrated analysis of metabolite and gene expression data. Here, metabolite data are considered the dependent (Y) variable and genes the explanatory variable. The tool allows for several combinations of metabolite and gene models. A note of caution: a complete metabolite and gene expression dataset with no filtering will be challenging to interpret using this tool.

We recommend that both gene expression and metabolite datasets be reduced to reflect a common biological hypothesis before running this tool. For example, metabolite data can be subset by class (i.e. using the 'Name_in_KEGG' column generated from the 'Link Name to KEGGID' tool). Users who want to include similarly behaving compounds without regard to identification or type of compound can estimate modules with the Modulated Modularity Clustering (MMC) tool (Stone and Ayroles 2009). Each module can be examined separately. Finally, metabolite data can be reduced by using both metabolite class and the MMC tool. Similarly, gene expression data can be reduced in scope by uploading a custom list of genes of interest or by using metagenes as implemented in PANA (Ponzoni et al. 2014).

1) Classes of metabolites can be modeled as a function of metagenes.
2) Classes of metabolites can be modeled as a function of a set of individual genes.
3) Unbiased clusters of metabolites can be modeled as a function of metagenes
4) Unbiased clusters of metabolites can be modeled as a function of a set of individual genes.

The tool executes a partial least squares regression with variable selection (sparse PLS, sPLS) as implemented in the 'mixOmics' package (Rohart F., Gautier, B, Singh, A and Lê Cao, K. A. mixOmics:

an R package for ‘omics feature selection and multiple data integration. On bioRxiv). The mixomics sPLS function is run in ‘classic mode’ (http://mixomics.org/methods/spls/) with the number of components included in the model set to 2. In addition, the user selects the number of variables (genes) for each component to use in model construction.

This tool needs at least 1 subset with a minimum number of 3 metabolites to run properly. If the user selects subset metabolites by class and no metabolite groups are identified or small metabolite groups with less than 3 members are found, the tool will stop and a warning message will be generated to try the MMC option instead. Similarly, if the user selects subset metabolites using MMC clusters and there are no clusters with at least 3 metabolites, the tool will stop and a warning message will be generated to try the 'by class’ option instead.

**Input:**

1. Select a **Metabolite Wide Dataset** for integration from your History.

2. Enter the name of the **column** in your Metabolite Wide Dataset that contains unique **FeatureIDs**.

3. **Optional**: Select a **Metabolite Annotation File.** Selecting this option allows the user to choose a column in the Annotation File for labeling output files so they are more user-friendly.

– if you select **'Yes'**

– Select **the Metabolite Annotation File** from your History.

– Enter the name of the **column** in your Metabolite Annotation File that you want to use to **label** the output files.

4. Select an option for reducing the Metabolite Data. You must select one of the options below:

**By metabolite class.** This will use a predefined grouping of metabolites.

All of the metabolites in each class (1,..j) will be treated as individual dependent variables in a multivariate regression ($Y_1$,..$Y_{n\_j}$) where j is the number of metabolites in class j. At least three metabolites in one class are needed for the tool to run.

-Select the **'Metabolite to KEGG ID Link**' File from your history. This file MUST contain a column called **'Name_in_KEGG'**. The names in this column are used to create the metabolite classes for analysis.

**By MMC pattern.** This will perform a MMC analysis using default parameters.

Each module will be analyzed as a set according to the MMC algorithm (Stone and Ayroles 2009). At least 3 metabolites in at least one module are needed.

-Select the '**Metabolite to KEGG ID Link**' File from your history.
-Select the **Design File** to use with your Metabolite KEGG ID Link File. This file can be generated using the 'Create: Design, Wide, and Annotation datasets' tool. Note that at minimum you need a column called 'sampleID' that contains the names of your samples.

-The next set of 4 options specify the **MMC running conditions**. We refer you to the MMC paper for details about what these options mean (Stone and Ayroles 2009).

-In the **Lower sigma value** text box, type the decimal lower sigma value from 0 to 1 range. The default value is 0.05.

-In the **High sigma value** text box, type the decimal upper sigma value from 0 to 1 range. The default value is 0.50. High sigma value has to be bigger than Lower sigma value for the algorithm to work.

-In the **Sigma values** text box, type the number of sigma values considered. The default is 451. Higher numbers increase the precision but decrease the performance time.

-Select the **Correlation method** from the drop-down method. (Pearson's correlation coefficient (default), Kendall's Tau correlation coefficient, Spearman's rank correlation).

**By both metabolite class and MMC pattern**. This will perform an MMC analysis using default parameters on a predefined grouping of metabolites.

All of the metabolites in each class (1,..j) will be treated as individual dependent variables in a multivariate regression ($Y_1$,..$Y_{n\_j}$) where j is the number of metabolites in class j. At least three metabolites in one class are needed for the tool to run.

-Select the '**Metabolite to KEGG ID Link**' File from your history. This file MUST contain a column called 'Name_in_KEGG'. The values in this column that are identical create the metabolite classes for analysis.

-Each module will be analyzed as a set according to the MMC algorithm (Stone and Ayroles 2009). At least 3 metabolites in at least one module are needed.

-Select the '**Metabolite to KEGG ID Link**' File from your history

-Select the **Design File** to use with your Metabolite KEGG ID Link File. This file can be generated using the 'Create: Design, Wide, and Annotation datasets' tool. Note that at minimum you need a column called 'sampleID' that contains the names of your samples.

The next set of 4 options specify the **MMC running conditions**. We refer you to the MMC paper for details about what these options mean (Stone and Ayroles 2009).

-In the **Lower sigma value** text box, type the decimal lower sigma value from 0 to 1 range. The default value is 0.05.

-In the **High sigma value** text box, type the decimal upper sigma value from 0 to 1 range. The default value is 0.50. High sigma value has to be bigger than Lower sigma value for the algorithm to work.

-In the **Sigma values** text box, type the number of sigma values considered. The default is 451. Higher numbers increase the precision but decrease the performance time.

-Select the **Correlation method** from the drop-down method. (Pearson's correlation coefficient (default), Kendall's Tau correlation coefficient, Spearman's rank correlation).

5. Select the **Gene Expression Wide Dataset** for integration from your History.

Enter the name of the **column** in your Gene Expression Wide Dataset that contains the unique **Feature IDs.**

Select whether you want to use a **Gene Expression Annotation File**. Selecting this option allows the user to choose a column in the Annotation File for labeling output files so they are more user-friendly.

– if you select **'Yes'**

– Select the **Gene Expression Annotation File** from your History.

– Enter the name of the **column** in your Annotation File that you want to use to **label** the output files.

6. Select which **option** to use for **reducing** the **Gene Expression Data**.

–Include **all genes** in the Gene Expression Wide Dataset -- all genes will be included in the analysis. Note: run times may be so long that the process fails. Not recommended.

– Upload a **custom lis**t of containing specific genes of interest - this list must be a single column containing the Gene Symbols for the genes of interest.

-Select a **Custom Gene List** from your history.

– Use **Metagenes** from PANA (Pathway network inference from gene expression data (Ponzoni, et al. 2014). Generates a gene expression submatrix per pathway (columns) by collecting expression profiles for the genes (rows) annotated to each pathway.

– Select the '**Gene to KEGG ID Link**' dataset from your history. This file can be generated using the 'Link Name to KEGGID' tool.

– Enter the name of the **column** in your 'Gene to KEGGID link' dataset that contains **Gene Symbols**

– Select the '**Gene KEGG Pathway File**' from your history. This file can be generated using the 'Add KEGG Pathway information' tool.

– Chose the **criterion to select components**. Details about these choices can be found in (Ponzoni et al. 2014).

1) **single%** = percentage of variability of that PC (default);

2) **%accum** = percentage of accumulated variability;

3) **abs.val** = absolute value of the variability of that PC;

4) **rel.abs** = fold variability of tot.var/rank(X)

– Enter **a Variability cut-off value**. Components with variability greater than this user-specified threshold value are kept in the model. Default value is 0.23. Details about these choices can be found in (Ponzoni et al. 2014).

**– Include Pathway Names in results files and figures**.

- If **'Yes'** then input the '**PathwayID2PathwayNames**' dataset. This dataset can be generated using the 'Add KEGG Pathway information' tool.

Given the above selections of what to model, the class or unbiased cluster for metabolites, the lists or metagenes from PANA, a sparse PLS is carried out using mixOmics (Rohart et al. 2016).

For the **sPLS**, the options below are required.

-Enter the **number of genes/metagene**s to keep for each component in the sPLS analysis. The default value is 10.

-Enter a **threshold value for returning correlations** in the sPLS output Files. Correlations less than this threshold will not be included in the output. This makes visualization of results manageable. Default value of 0.8.

**Output:**

For metabolite reduction by metabolite class and all genes:

**(1) A PDF containing a sPLS figure for each metabolite class.** In the sphinogomyelin example shown below, gene symbols are on the Y-axis and metabolites are on the X-axis.

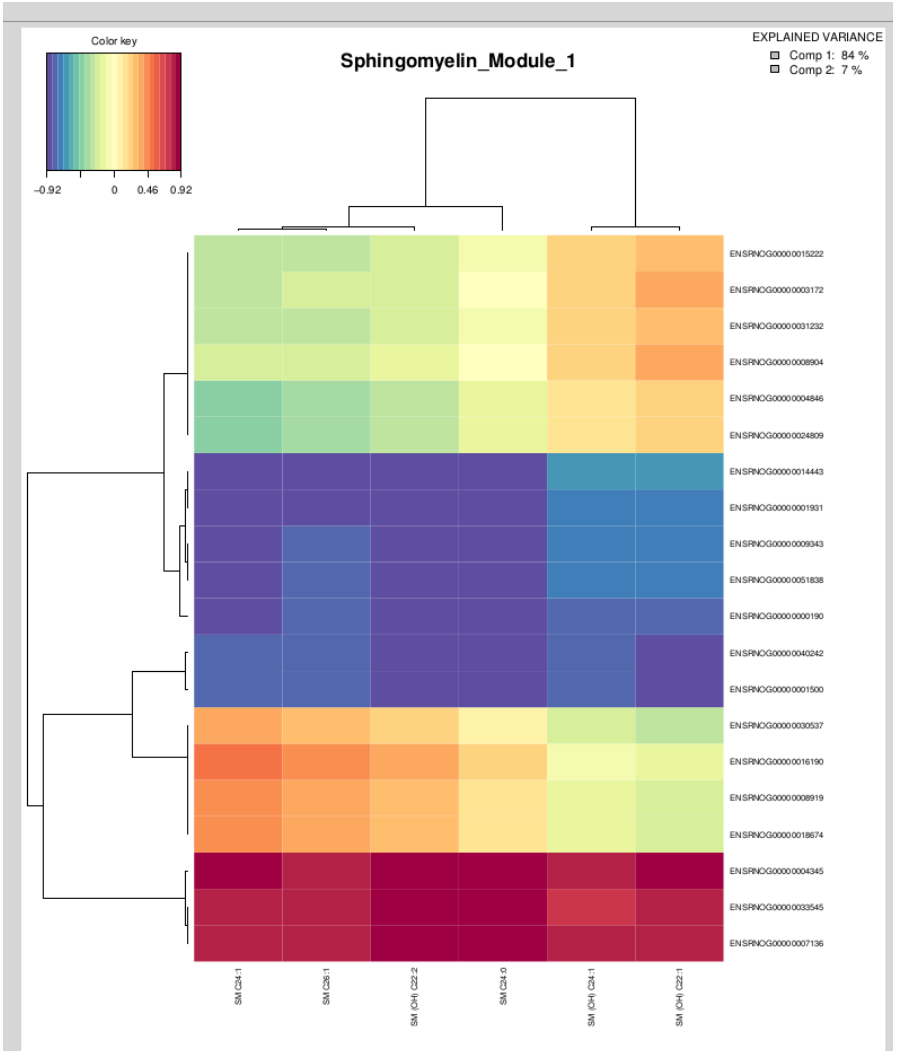


(2) A **sPLS Correlation TSV file** containing the correlations for each metabolite-gene pair and what metabolite class (subset) the pair locate to.

Analyze Data Workflow

| 1 | 2 | 3 | 4 |
|---|---|---|---|
| Metabolite | Gene | Correlation | Subset |
| Met_6 | Gene_11759 | 0.952 | Sphingomyelin |
| Met_6 | Gene_6035 | -0.919 | Sphingomyelin |
| Met_11 | Gene_2899 | -0.906 | Sphingomyelin |
| Met_6 | Gene_7684 | 0.906 | Sphingomyelin |
| Met_11 | Gene_11862 | -0.903 | Sphingomyelin |
| Met_11 | Gene_10889 | 0.899 | Sphingomyelin |
| Met_11 | Gene_3598 | -0.897 | Sphingomyelin |
| Met_3 | Gene_11862 | -0.895 | Sphingomyelin |
| Met_3 | Gene_3598 | -0.889 | Sphingomyelin |
| Met_6 | Gene_6567 | 0.889 | Sphingomyelin |
| Met_6 | Gene_2899 | 0.887 | Sphingomyelin |
| Met_3 | Gene_2899 | -0.887 | Sphingomyelin |
| Met_3 | Gene_10889 | 0.886 | Sphingomyelin |
| Met_11 | Gene_1454 | -0.885 | Sphingomyelin |
| Met_11 | Gene_2002 | -0.876 | Sphingomyelin |
| Met_3 | Gene_1454 | -0.873 | Sphingomyelin |
| Met_11 | Gene_6567 | -0.873 | Sphingomyelin |
| Met_8 | Gene_11759 | 0.868 | Sphingomyelin |

**For metabolite reduction by MMC:**

If the user selects the option to subset metabolites using MMC, the outputs include, in addition to the **sPLS PDF** and the **sPLS Correlation TSV file** described above:

(3) **A MMC PDF Figure containing unsorted, sorted, and sorted-smoothed heatmaps of the variance- covariance matrixes.**

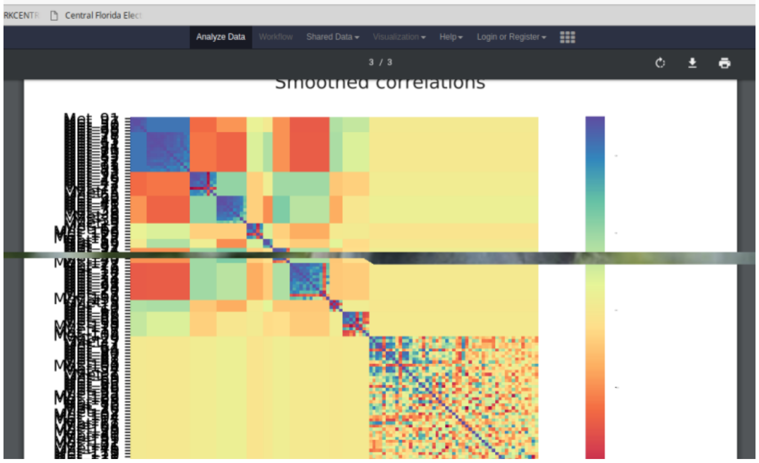


(4) **A MMC Output TSV file** containing algorithm summaries in the following columns:

**Unique metabolite featureID**

**Module**: Contains the module number for each feature calculated by the MMC tool.

**Entry Index**: Contains the original order of the names of the rows of the input Metabolite Wide Dataset.

**Degree**: Average of the absolute values of correlations for the given element in a block to other elements within that block.

**Average Degree**: Average values of the degrees computed above across all elements within the given block.

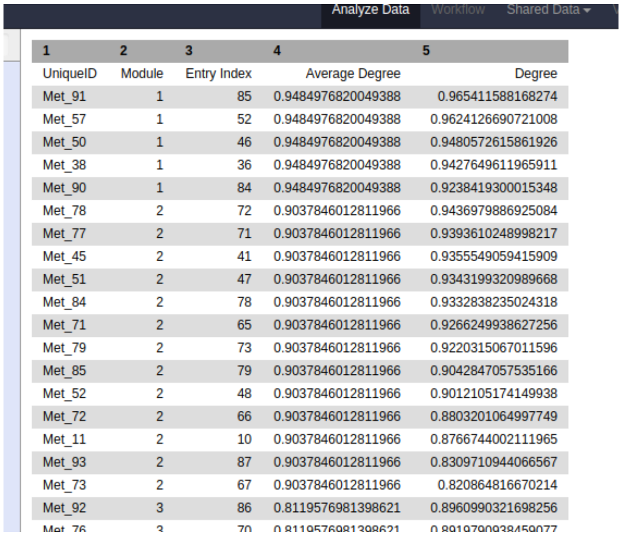


| 1 | 2 | 3 | 4 | 5 |
|---|---|---|---|---|
| UniqueID | Module | Entry Index | Average Degree | Degree |
| Met_91 | 1 | 85 | 0.9484976820049388 | 0.965411588168274 |
| Met_57 | 1 | 52 | 0.9484976820049388 | 0.9624126690721008 |
| Met_50 | 1 | 46 | 0.9484976820049388 | 0.9480572615861926 |
| Met_38 | 1 | 36 | 0.9484976820049388 | 0.9427649611965911 |
| Met_90 | 1 | 84 | 0.9484976820049388 | 0.9238419300015348 |
| Met_78 | 2 | 72 | 0.9037846012811966 | 0.9436979886925084 |
| Met_77 | 2 | 71 | 0.9037846012811966 | 0.9393610248998217 |
| Met_45 | 2 | 41 | 0.9037846012811966 | 0.9355549059415909 |
| Met_51 | 2 | 47 | 0.9037846012811966 | 0.9343199320989668 |
| Met_84 | 2 | 78 | 0.9037846012811966 | 0.9332838235024318 |
| Met_71 | 2 | 65 | 0.9037846012811966 | 0.9266249938627256 |
| Met_79 | 2 | 73 | 0.9037846012811966 | 0.9220315067011596 |
| Met_85 | 2 | 79 | 0.9037846012811966 | 0.9042847057535166 |
| Met_52 | 2 | 48 | 0.9037846012811966 | 0.9012105174149938 |
| Met_72 | 2 | 66 | 0.9037846012811966 | 0.880320106497749 |
| Met_11 | 2 | 10 | 0.9037846012811966 | 0.876674400211965 |
| Met_93 | 2 | 87 | 0.9037846012811966 | 0.8309710944066567 |
| Met_73 | 2 | 67 | 0.9037846012811966 | 0.820864816670214 |
| Met_92 | 3 | 86 | 0.8119576981398621 | 0.8960990321698256 |
| Met_76 | 3 | 70 | 0.8119576981398621 | 0.8919790938459077 |

**For subsetting genes by generating metagenes using PANA:**

If the user selects the option to subset genes using PANA, the outputs include, in addition to the **sPLS PDF** and **the sPLS Correlation TSV file** above:

(5) **A PANA Output TSV table** containing associations between gene symbols and KEGG pathways.

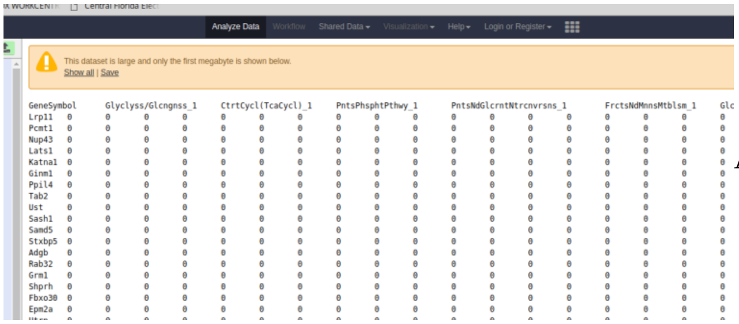


| GeneSymbol | | Glyclyss/Glcngnss_1 | | | CtrtCycl(TcaCycl)_1 | | | PntsPhsphtPthwy_1 | | | PntsNdGlcrntNtrcnvrsns_1 | | | | FrctsNdMnnsMtblsm_1 | | | Glc |
|---|---|---|---|---|---|---|---|---|---|---|---|---|---|---|---|---|---|---|
| Lrp11 | 0 | 0 | 0 | 0 | 0 | 0 | 0 | 0 | 0 | 0 | 0 | 0 | 0 | 0 | 0 | 0 | 0 | 0 |
| Pcmt1 | 0 | 0 | 0 | 0 | 0 | 0 | 0 | 0 | 0 | 0 | 0 | 0 | 0 | 0 | 0 | 0 | 0 | 0 |
| Nup43 | 0 | 0 | 0 | 0 | 0 | 0 | 0 | 0 | 0 | 0 | 0 | 0 | 0 | 0 | 0 | 0 | 0 | 0 |
| Lats1 | 0 | 0 | 0 | 0 | 0 | 0 | 0 | 0 | 0 | 0 | 0 | 0 | 0 | 0 | 0 | 0 | 0 | 0 |
| Katna1 | 0 | 0 | 0 | 0 | 0 | 0 | 0 | 0 | 0 | 0 | 0 | 0 | 0 | 0 | 0 | 0 | 0 | 0 |
| Ginm1 | 0 | 0 | 0 | 0 | 0 | 0 | 0 | 0 | 0 | 0 | 0 | 0 | 0 | 0 | 0 | 0 | 0 | 0 |
| Ppil4 | 0 | 0 | 0 | 0 | 0 | 0 | 0 | 0 | 0 | 0 | 0 | 0 | 0 | 0 | 0 | 0 | 0 | 0 |
| Tab2 | 0 | 0 | 0 | 0 | 0 | 0 | 0 | 0 | 0 | 0 | 0 | 0 | 0 | 0 | 0 | 0 | 0 | 0 |
| Ust | 0 | 0 | 0 | 0 | 0 | 0 | 0 | 0 | 0 | 0 | 0 | 0 | 0 | 0 | 0 | 0 | 0 | 0 |
| Sash1 | 0 | 0 | 0 | 0 | 0 | 0 | 0 | 0 | 0 | 0 | 0 | 0 | 0 | 0 | 0 | 0 | 0 | 0 |
| Samd5 | 0 | 0 | 0 | 0 | 0 | 0 | 0 | 0 | 0 | 0 | 0 | 0 | 0 | 0 | 0 | 0 | 0 | 0 |
| Stxbp5 | 0 | 0 | 0 | 0 | 0 | 0 | 0 | 0 | 0 | 0 | 0 | 0 | 0 | 0 | 0 | 0 | 0 | 0 |
| Adgb | 0 | 0 | 0 | 0 | 0 | 0 | 0 | 0 | 0 | 0 | 0 | 0 | 0 | 0 | 0 | 0 | 0 | 0 |
| Rab32 | 0 | 0 | 0 | 0 | 0 | 0 | 0 | 0 | 0 | 0 | 0 | 0 | 0 | 0 | 0 | 0 | 0 | 0 |
| Grm1 | 0 | 0 | 0 | 0 | 0 | 0 | 0 | 0 | 0 | 0 | 0 | 0 | 0 | 0 | 0 | 0 | 0 | 0 |
| Shprh | 0 | 0 | 0 | 0 | 0 | 0 | 0 | 0 | 0 | 0 | 0 | 0 | 0 | 0 | 0 | 0 | 0 | 0 |
| Fbxo30 | 0 | 0 | 0 | 0 | 0 | 0 | 0 | 0 | 0 | 0 | 0 | 0 | 0 | 0 | 0 | 0 | 0 | 0 |
| Epm2a | 0 | 0 | 0 | 0 | 0 | 0 | 0 | 0 | 0 | 0 | 0 | 0 | 0 | 0 | 0 | 0 | 0 | 0 |

ADDIN EN.REFLISTX


Ponzoni, I., M. J. Nueda, S. Tarazona, S. Gotz, D. Montaner, J. S. Dussaut, J. Dopazo & A. Conesa

(2014) Pathway network inference from gene expression data. *Bmc Systems Biology*.

Rohart, F., B. Gautier, A. Singh & L. C. K-A. 2016. mixOmics: An R package for 'omics feature selection and multiple data integration.

Stone, E. A. & J. F. Ayroles (2009) Modulated Modularity Clustering as an Exploratory Tool for Functional Genomic Inference. *Plos Genetics*, 5.

Wu, C. L., I. MacLeod & A. I. Su (2013) BioGPS and MyGene.info: organizing online, gene-centric information. *Nucleic Acids Research*, 41**,** D561-D565.
